# A detailed investigation of Shared Variance Component Analysis as a tool to characterize neural dimensionality

**DOI:** 10.64898/2026.04.30.721904

**Authors:** Alejandro Carballosa, Alessandro Torcini

## Abstract

**Background:** The relevance of spontaneous activity has been unlocked thanks to recent large scale recordings that revealed, via Shared Variance Component Analysis (SVCA), the high-dimensional nature of the ongoing activity. A fundamental problem is how the dimension modifies when more neurons are included in the analysis. Contradictory results have been reported on this subject based on SVCA and Principal Component Analysis (PCA).

**New Method:** We investigate *pro et contra* of SVCA and PCA for the identification of reliable responses encoding underlying state variables. We focus on common features of the spectra of the reliable variances (RVs) and on their dimensionality. The analysis is demonstrated on previously published Ca2^+^ data from the visual and the dorsal cortex in head fixed mice during spontaneous behavior.

**Results:** RVs grow proportionally to the number *N* of neurons and show a power-law decay *k*^−*α*^ with the *k*-th SVC dimension over a range bounded by a maximal dimension *k*_*c*_, initially diverging as *N* ^1*/α*^ and then saturating at sufficiently large *N*. The reliable dimensionality, estimated with different methodologies, also shows a clear saturation to an asymptotic value for large *N*. Furthermore, its value decreases when *α* becomes larger, as demonstrated by employing experimental data as well as theoretical predictions.

**Conclusion:** We have shown that SVCA is an extremely effective tool to extract reliable features from the neural signals, and that the exponent *α* represents a biomarker able to reveal the level of correlation of the neurons as well as the dimensionality of the reliable space.

**Highlights:**

- Advantages and drawbacks of Shared Variance Component Analysis to extract reliable signals from neural data
- Comparison of different methods to estimate reliable neural dimensionality associated to spontaneous activity
- Analytical expressions of embedding dimensionality for power-law decaying reliable variances
- Bounded growth of the dimensionality with the number of neurons

## 1. Introduction

As neural recordings have grown to encompass thousands of simultaneously recorded neurons, dimensionality reduction has become a foundational approach in contemporary systems neuroscience (Urai et al., 2022; Stringer and Pachitariu, 2024). These techniques facilitate the exploration and interpretation of large-scale data sets and they can be classified into two main classes according to which type of dimensionality they capture (see (Mitchell-Heggs et al., 2023) for a detailed review).

Following (Jazayeri and Ostojic, 2021), we can distinguish between *intrinsic* and *embedding* dimensionality. The intrinsic dimension is based on non-linear data analysis and gives the minimal number of degrees of freedom (independent variables) needed to characterize the evolution on the attractor, this is also usually referred as attractor dimension (Grassberger and Procaccia, 1983; Kantz and Schreiber, 2003). The embedding dimension instead gives a measure of the number of relevant dimensions covered by the attractor (manifold) within the original phase space, and its estimation is usually based on linear transformations of the original data, as those employed in Principal Component Analysis (PCA) (Sirovich, 1989) and Shared Variance Component Analysis (SVCA) (Stringer et al., 2019b). More specifically, PC-based methodologies decompose the data into linear combinations of neural population vectors (components) and evaluate the embedding dimension by estimating the number of components required to explain most of the data’s total variance. These methods are widely used for their computational simplicity and easy biological interpretation (Cunningham and Yu, 2014; Mazzucato et al., 2016; Mitchell-Heggs et al., 2023), with the most relevant components (the ones associated to larger variances) often correlating with behavior, task, or stimuli (Gao et al., 2017). Methods based on linear decomposition have been important in revealing the underlying neural mechanisms behind motor tasks and control (Churchland et al., 2012; Kaufman et al., 2014; Gallego et al., 2017), decision-making in the prefrontal cortex (Mante et al., 2013; Harvey et al., 2012) or working memory (Cueva et al., 2020), among others (see (Urai et al., 2022; Stringer and Pachitariu, 2024) for a reviews on the subject). On the other hand, non-linear methods, such as uniform manifold approximation and projection (UMAP) or t-distributed stochastic neighbor embedding (tSNE), attempt to approximate the topology of the population dynamics’ attractor and to map it onto a given low-dimensional space (Stringer and Pachitariu, 2024). These methods are particularly suited to analyze complex tasks that might generate highly curved, nonlinear manifolds associated to the neural dynamics (Jazayeri and Ostojic, 2021). While nonlinear methods are more suited to reveal the intrinsic dimensionality of the data, their biological interpretability is often weaker, as the recovered manifolds are not always easily decodable. Moreover, they might not always yield a good reconstruction of the original data if these are high dimensional (Torcini et al., 1991; Mitchell-Heggs et al., 2023). For this reason, nonlinear methods are frequently combined with an initial linear preprocessing to extract the most salient features of the data. Such hybrid approaches have successfully uncovered neural representations in head direction cells (Chaudhuri et al., 2019) and grid cells (Gardner et al., 2022), and revealed that the topology of neural activity often mirrors that of the task or environment (Yang et al., 2024). However, since our study focuses on spontaneous neural activity in large-scale data sets in absence of any task or stimuli, we will limit to consider linear embedding dimensions.

An important question, yet still open, is how the estimated dimensionality scales with the number *N* of recorded neurons (Gao and Ganguli, 2015). This question has been previously addressed by Mazzucato et al. (2016). For homogeneous networks, where all neurons exhibit the same pair-wise correlation *ρ*, these authors provided a theoretical upper bound for the dimensionality to which the dimension should saturate for sufficiently large *N*. Moreover, for independent neurons (with *ρ* = 0) they show that the dimensionality should grow unbounded and proportionally to *N*, analogously to what was recently observed in balanced recurrent neural networks (Clark et al., 2023; Engelken et al., 2023). More recent PC-based analysis confirm the saturation of the linear embedding dimensions for different type of neural data (obtained via neuropixel probes and calcium imaging) as well as for different cortical areas in spontaneous behaving mice (Dahmen et al., 2020) and for the whole brain of larval zebrafish hunting and in spontaneous conditions (Wang et al., 2025). Complementarily, Williamson et al. (2016) used factor analysis (FA) to distinguish shared dimensionality (i.e. the number of modes with shared covariability) from shared variance across neurons for a given mode (i.e. the fraction of total variance shared across neurons for that dimension). Analyzing subsets of *N* = 20 − 80 neurons, they found that shared dimensionality increased with *N*, whereas the total shared variance and the most dominant modes remained stable. This indicates that recording more neurons reveals additional shared dimensions, but that these new dimensions account for little extra shared variance, since the total remains essentially unchanged. All together, these works suggest that population encoding is performed redundantly across the neural population, which implies that large and detailed data sets might not be necessary to decode the main features of the population dynamics.

On another side, recent experimental advances enabling large-scale simultaneous recordings challenged this notion by revealing that the dimensionality of neural representations, during spontaneous activity, has been likely underestimated because of a low sampling of neurons, finding high-dimensionality structure in the data that can span hundreds of linear dimensions (Stringer et al., 2019b) or even display an *unbounded* growth of the measured dimensionality with the number of neurons up to one million neurons (Manley et al., 2024b). Based on these results, Stringer et al. (2019b) and Manley et al. (2024b) propose that larger populations would capture an expanding set of internal variables distributed across the whole cortex and that cannot be explained only by arousal and behavioral states. However, the origin, role and relevance of these internally generated variables remain as unanswered questions (Avitan and Stringer, 2022). These studies are based on SVCA, which represents an important refinement of PCA for spontaneous activity by including a cross-validation of the population activity across neurons and timepoints. This cross-validation allows for the identification of reliable components (Shared Variance Components (SVCs)), which still represent directions along which neurons maximally covary in time, but removing sources of noise and erraticity intrinsic to the dynamics of each single neuron. Reliable signals are present during spontaneous neural activity and are due to the encoding of common unobservable state variables. The cross-validation appears to be the key ingredient to unravel power-law decay of the reliable variances with the SVC dimensions in spontaneous (Stringer et al., 2019b; Manley et al., 2024b) and stimulus-evoked activity (Stringer et al., 2019a). In particular, a relationship between the measured power-law exponents and the data smoothness and dimensionality have been suggested in (Stringer et al., 2019a).

In order to somehow reconciliate these two apparently contraddictory visions, in this paper we will re-examine in details the SVCA approach to identify advantages and drawbacks with respect to standard PCA. In particular, we have considered synthetic data as well as openly available large neural data sets relative to spontaneous activity in awake, head fixed mice measured with calcium imaging (Stringer et al., 2019b; Syeda et al., 2024; Manley et al., 2024b).

Synthetic data allows for the detailed characterization of SVC reliable spectra and identification of possible scaling laws with the number of neurons as well as for the identification of drawbacks of the methodology. Following this preliminar analysis we have then considered the power-law decaying reliable variances (spectra) associated to the finite neural data sets and derived their asymptotic profile for an infinite number of neurons. Moreover, we have estimated SVC-based embedding dimensionality for the considered data sets and compared the experimental findings with analytical expressions for such dimensionality for finite and infinite number of neurons. As a last aspect, we have examined PC-based embedding dimensions for the same data sets and compared the results to SVC-based estimations.

The paper is structured as follows. In section 2 we review briefly PCA and the SVCA in more details, we then report methods usually employed to measure PC-based emedding dimensionality and discuss their extention to SVCA. Furthermore, in the same section we describe the synthetic and experimental data analyzed in the remaining of the paper. The section 3 is divided in two main subsections: namely, subsections 3.1 and 3.2. In the first one (subsection 3.1) we consider synthetic noisy data to understand the ability and the drawbacks of the SVCA, as compared to PCA, to recover the reliable responses induced by the common underlying signals subject to noise. In the subsection 3.2 we will apply PCA and SVCA to neural data sets corresponding to spontaneous neural activity. In particular, for these data we will identify the main common features of the reliable and total SVC spectra and characterize the dependence of the reliable and total variances on the number of considered neurons. We will then focus on the estimation of dimensionality of the data sets by employing different SVC-based indicators to estimate the reliable embedding dimensions and PC-based indicator to extract the dimensionality of the whole linear space. Finally in section 4 we summarize our findings and discuss their relevance in the context of the existing literature.

## 2. Methods

### 2.1. Principal Component and Shared Variance Component Analyses

In this subsection we will give a brief overview of the well known PCA or Karhunen-Loève decomposition (Sirovich, 1989) and of the more recently introduced SVCA (Stringer et al., 2019b).

#### 2.1.1. Principal Component Analysis

For a given data set, PCA corresponds to the eigen-decomposition of the covariance matrix which provides eigenvalue and eigenvectors associated to the considered matrix. In particular, PCA provides the orthogonal directions (eigenvectors) and the associated fraction of variance explained (eigenvalue). The number of principal components (PCs) necessary to reconstruct the majority of the total variance gives an estimate of the number of degrees of freedom of the underlying dynamics. If large part of the variance can be obtained with few PCs, the second-order statistics of the collective dynamics is constrained in a hyper-ellipsoid with few axes.

More in detail, let **X** be a (*N* × *T*) matrix representing the neural activity {*x*_*i*_(*t*)} *i* = 1, …, *N* of *N* neurons registered at regular time intervals Δ*t* for *T* successive time steps. PCA transforms the data matrix **X** linearly by rotating it onto a set of orthogonal, uncorrelated dimensions where covariance is maximized. In other words, its purpose is to find a good representation of **X** in a set of variables at lower dimensionality where redundancy - given by the correlations in the data structure - is removed (Hyvärinen et al., 2001).

One standard pre-processing step in PCA is doing a z-scoring transformation (substraction of the mean and division by the standard deviation) of the data to ensure that all variables contribute equally to the variance of the signal. This process also ensures that the cross-correlation matrix equals the *N* × *N* covariance matrix **K** that can be directly computed as

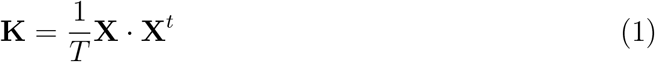

where each element of the matrix 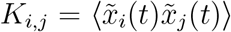 epresents the temporal average ⟨·⟩ of the product of the z-scored neural activity 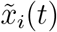 of neuron *i* and of that of neuron 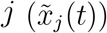 taken at the same time *t*. The PCs of the data **X** are the eigenvectors 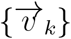 of the covariance matrix **K**, in other words we can express the neural activity on this new orthonormal basis, as

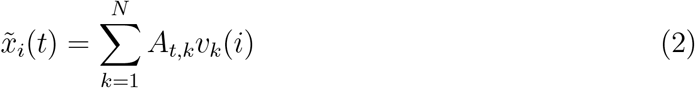

where *v*_*k*_(*i*) is the *i*-th component of the *k*-th eigenvector, and the corresponding non-negative eigenvalue *λ*_*k*_ is given by

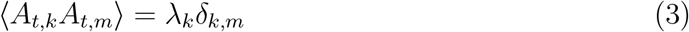

where *δ*_*k,m*_ is the Kroenecker *δ*-function. Therefore *λ*_*k*_ represents the average squared size (variance) of the data set along the direction 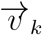.

#### 2.1.2. Shared Variance Component Analysis

Recently, Stringer et al. (2019b) have introduced a variation of the PCA, termed SVCA, with the aim to reveal components (dimensions) of the neural variance associated with the reliable responses induced by common underlying signals. In particular, SVCA methodology has been specifically developed to analyze single-trial, spontaneous neuronal activity.

The main idea behind the method is that spontaneous neural activity can be split in two components: namely, a reliable one which represents the deterministic encoding of an unobservable state variable and an unreliable one, which is independent among the neurons and somehow erratic. For example, the reliable component can be the mean population firing rate, which can be related to some hidden behavioural state variable, and the unreliable ones the chaotic or stochastic fluctuations of the single neuron firing rates.

First of all, Stringer et al. (2019b) assumed schematically that the activity *f*_*k*_(*t*) of the neuron *k* at time *t* can be written as the sum of a reliable signal *F*_*k*_(*s*_*t*_) that depends deterministically on a state variable *s*_*t*_ at time *t* plus a noise term *ξ*_*k*_(*t*) (with zero average and variance 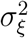) that is independent among neurons and of the state *s*_*t*_:

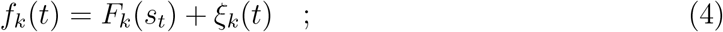

the state variable *s*_*t*_ can take *S* possible values distributed accordingly to some probablilty distribution function (PDF) *P* (*s*). In order to estimate the reliable components of the signal the neurons are split in two populations of size *N* and *M*, with corresponding activity vectors *f* (*t*) ∈ ℝ^*N*^ and *g*(*t*) ∈ ℝ^*M*^, and the recorded activity of the neurons is divided in two equal halves made of *T* timepoints, termed *training* and *test* sets. The training set is used to identify the directions along which the neural activity of the two population of neurons present the maximal covariance, these are the shared variance components (SVCs). The activity of the whole population of neurons during the test period is then projected into each SVC-vector, previously identified in the training period. An estimate of the *reliable variance* (RV) along a certain direction SVC is then obtained by estimating the correlation between the projections from the two neural populations.

In practice, by employing the training data sets one defines a (*N* × *M*) cross-covariance matrix **C**, whose elements are given by

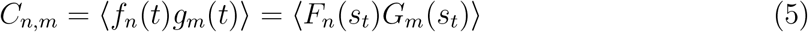

the averages ⟨·⟩ are time averages over the *T* training points. By assuming that the noise terms have average zero and are independent one gets

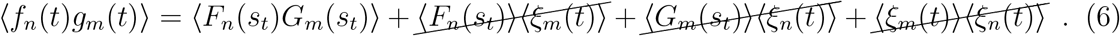

Finally the time average ⟨*F*_*n*_(*s*_*t*_)*G*_*m*_(*s*_*t*_)⟩ would be equivalent to an average over all the possible states *s*_*t*_ if *T* is sufficently long and the dynamics is ergodic, namely

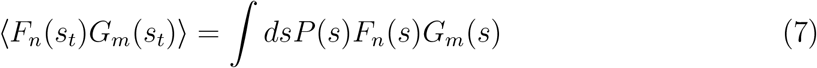

Note that for the PCA, where the covariance is estimated among the same set of neurons, the terms of the covariance matrix will read as

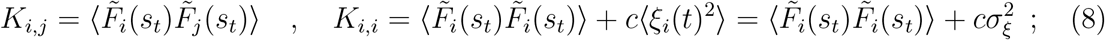

where *c* is some normalization factor taking in account the z-scoring of the data. Therefore the diagonal terms of the covariance matrix are strongly affected by the noise terms and the variance of the noise 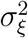 will clearly contribute to the variances {*λ*_*i*_} along the principal components.

The cross-covariance matrix can be formally written as 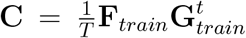, where **F**_*train*_ (**G**_*train*_) is a *N* × *T* (*M* × *T*) matrix representing the z-scored neural activity 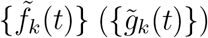 during the training period of the first (second) population of *N* (*M*) neurons. The singular value decomposition of the matrix **C** takes the following expression

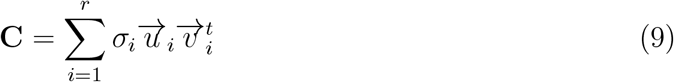

where 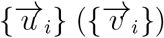 are orthonormal basis in ℝ^*N*^ (ℝ^*M*^) termed left (right) singular vectors of **C**, while *σ*_*i*_ are real non increasing values with *r ≤* Min {*M, N*}.

An estimate of the RV associated to the *k*-th SVC is obtained by projecting the activity of the two populations during the *test* period along the *k*-th left and right singular vectors (obtained during the *training* period) and by estimating their cross-correlation, as follows

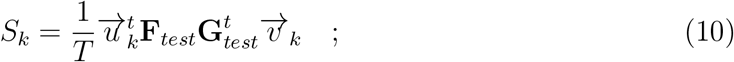

where **F**_*test*_ (**G**_*test*_) is a *N* × *T* (*M* × *T*) matrix containing the neural activity during the test phase of the first (second) neural population.

This quantity *S*_*k*_ should be compared with the *total variance* (TV) *S*_*k,tot*_ along the direction *k*, that can be estimated as the mean of the variances of the test data set projected along the *k*-th right and left singular vectors obtained for the training set, namely

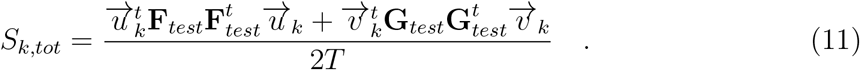

The ratio of these two quantities

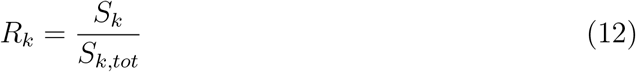

gives the fraction of reliable variance contained in the *k*-th SVC, therefore the fraction of the variance associated to independent *noise* along this direction is 1 − *R*_*k*_. For brevity, we will term the ensemble of the {*S*_*k*_} ({*S*_*k,tot*_}) values, ordered from the largest to the smallest, the *reliable* (*total*) SVC spectrum, even if the quantities *S*_*k*_ (*S*_*k,tot*_) are not proper eigenvalues nor singular values on the consider basis.

Indeed, the *k*-dimensional reliable variance 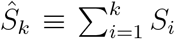 obtained from the projection of the test time-points does not necessarily coincide with the actual variance of the matrix **F**_*test*_**G**_*test*_, but it rather expresses how the variance of the latter matrix falls in the directions of maximal covariance of the training time-points associate to the matrix **F**_*train*_**G**_*train*_. For this reason, *Ŝ*_*k*_ is a lower bound for the sum of the largest *k* singular values 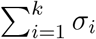 of the matrix **C**, that *Ŝ*_*k*_ approaches when the number of training time-points becomes sufficiently large. Stringer et al. (2019b) have also proofed that if the number of neurons is sufficiently large, 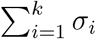 converges to the reliable variance of the entire neural population’s top *k* SVCs.

In the following we will show explicitly that because of this conceptual difference, the *reliable* SVC spectrum {*S*_*k*_} can contain negative values that become larger as the eigenvectors of the training data sets differ from those of the test data sets. In particular, we will show that these negative RVs decay with the number of time-points *T* in the test data set as *T*^−1*/*2^. To perform such analysis we will consider the sum of all the positive (negative) RVs defined as *Ŝ*^+^ (*Ŝ*^−^).

For simplicity, we have always set *N* = *M* and called this the system size, even if the total number of considered neurons for the SVC procedure is indeed 2*N* and not *N* as for the PCA.

#### 2.1.3. Estimation of the embedding dimensionality of the neural data

The activity (firing rate) of a sample of *N* neurons can be seen as a point evolving in an *N* dimensional phase space, with the time evolution of the population activity usually being restricted to an attractor (manifold) of dimension strictly smaller than *N*. It is important to estimate the value of this dimension since it will give a measure of the complexity of the neural activity and of the number of variables needed to describe such evolution. As already mentioned in the Introduction, one could characterize the neural activity in terms of the non-linear *intrinsic* (or attractor) dimension and of the linear *embedding* dimension (Jazayeri and Ostojic, 2021). Recently the embedding and intrinsic dimensions have been compared in (Engelken et al., 2023) for recurrent neural networks, revealing that the PC-based embedding dimension, measured via participation ratio, can give a reasonable estimate of the attractor dimension, measured via the Lyapunov (or Kaplan-Yorke) dimension (Kantz and Schreiber, 2003). However, the participation ratio does not represent an upper or lower bound for the attractor dimension, since it can be either larger or smaller (Engelken et al., 2023). In this paper we will limit to consider linear embedding dimensions for the analysis of neural data.

In particular, for what concerns PCA in order to give an estimation of the embedding dimension, it is customary to order the eigenvalues from the largest to the smallest and to introduce the cumulative variance 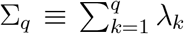. The minimal number *D*_*PCA*_(*p*) of the PC eigenvalues necessary to recover an arbitrary percentage *p* of the whole variance Σ_*N*_, i.e.

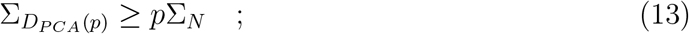

will give an estimate of the linear notion of the system dimensionality. In particular, *D*_*PCA*_(*p*) is the dimension of the linear subspace of the principal components containing the percentage *p* of the variance of the whole data set. Usually the ratio Σ_*k*_*/*Σ_*N*_ is reported as a function of the considered eigenvector *k* to assess the percentage of recovered variance with *k* PC eigenmodes.

Another estimation of the embedding dimensionality, always based on the PC eigen-values, but not requiring the introduction of an arbitrary threshold of variance, can be obtained by employing the so-called *participation ratio* (Gao et al. (2017))

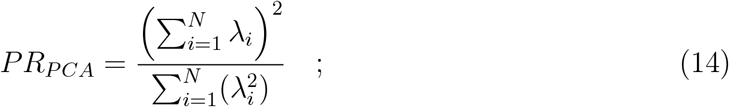

which has been recently employed to characterize the activity of chaotic neural networks in terms of PCA (Engelken et al., 2023; Clark et al., 2023; Kuśmierz et al., 2025) and even extended to take into account scale-dependent features (Recanatesi et al., 2022). This quantity gives an estimation of how many dimensions of the PC eigenspectrum are relevant, relative to the total variance and it has been suggested to correspond to *D*_*PCA*_(*p*) with *p* = 80% − 90% (Gao et al., 2017).

It is important for the following analysis to focalize on the characterization of a pure white noise signal. In particular, if the activity of the *N* neurons is generated as Gaussian noise with zero mean and unitary variance, we expect that, for time-series of sufficiently long duration *T*, the PC eigenvalues will become all identical *λ*_*i*_ = 1 ∀*i* = 1, …, *N*, therefore the embedding dimension will coincide with that of the whole the phase space, i.e. *D*_PCA_(*p*) = *PR*_PCA_ = *N*, since Σ_*N*_ = *N* for *z*-scored variables.

In this paper we will employ the previously reported methods (13) and (14), also in the context of the SVCA, to give an estimation of the dimensionality of the reliable components present in the neural dynamics. In particular this will amount to substitute the PC eigenvalues {*λ*_*i*_} appearing in (13) and (14) with the RVs {*S*_*k*_} and the cumulative PC variances Σ_*q*_ with *Ŝ*_*q*_. The corresponding dimensions will be referrred as *D*_*SVCA*_ and *PR*_*SVCA*_. These quantities will be estimated by limiting the sums appearing in (13) and (14) only to positive *S*_*k*_, whenever negative values appear in the reliable SVC spectrum.

### 2.2. Mathematical Model: the Lorenz Model

The Lorenz model (Lorenz, 1963a) is a simplified non-linear model for Rayleigh–Benard convection in two spatial dimensions limited to three spatial Fourier modes representing the main features of the convective motion. This model represented the first paradigmatic example of chaotic dynamics in a continuous low-dimensional deterministic system. The model is written as follows:

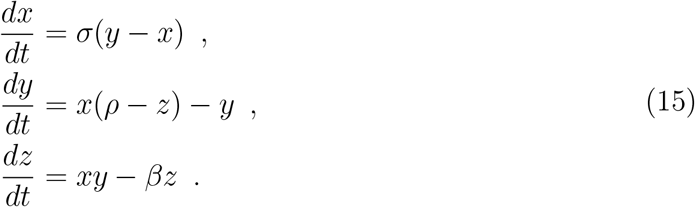

where *x*(*t*) is the amplitude of the convective motion, *y*(*t*) the temperature difference between ascending and descending fluid, and *z*(*t*) the deviation from the linear temperature profile. The parameters (*σ, ρ, β*) have physical meaning linked to the characteristic of the fluid, to the temperature and to the geometric shape of the convection cell. The parameters are selected to observe a chaotic dynamics: namely, *σ* = 10, *ρ* = 28, *β* = 8*/*3. In this case the intrinsic (correlation) dimension is well known and takes the value *D*_*f*_ = 2.05 ± 0.01 Grassberger and Procaccia (1983). This amounts to have [*D*_*f*_ + 1] active degrees of freedom (latent independent variables) in a 3-dimensional phase space, where [·] represents the entire part of a real number.

We have simulated the system (15) with a a 4th order Runge-Kutta scheme with time step Δ*t* = 0.01 for a time span *T* Δ*t* and registered the data corresponding to the evolution of the coordinates (*x*(*t*), *y*(*t*), *z*(*t*)) in a 3 × *T* matrix **X**^*′*^.

To prepare the data to be analysed in the following with PC and SVC methods, we *lift* the 3 dimensional space to an *N* dimensional space by performing a linear transformation. In particular, we construct a projection matrix **P** ∈ ℝ^3*×N*^ with entries sampled independently from the normal distribution 𝒩 (0, 1), where we set *N* = 1000.

The lifting of the Lorenz dynamics into the *N*-dimensional space is obtained by performing the following linear transformation

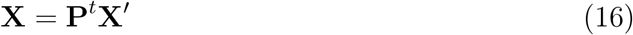

where **X** ∈ ℝ^*N ×T*^ contains the lifted coordinates at each time step.

#### 2.2.1. Description of the noise sources

In order to test the robustness of the methods against noise, we will add to the known deterministic dynamics different types of noise *ξ*(*t*).

In particular, we have firstly considered uncorrelated white Gaussian noise *ξ*(*t*) ∈ 𝒩 (0, 𝒞^2^) with zero mean and standard deviation 𝒞. The latter is related to the amplitude of the deterministic signal *A*(*t*) by the noise-to-signal ratio *ϵ*,

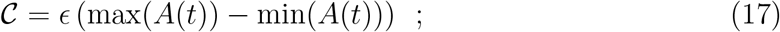

where the maximum and minimum are taken on the time interval during which noise is applied.

Since in neural systems noise sources are usually far from being uncorrelated, we have also considered colored noise modeled through an Ornstein - Uhlenbeck (OU) process (San Miguel and Toral (2000)),

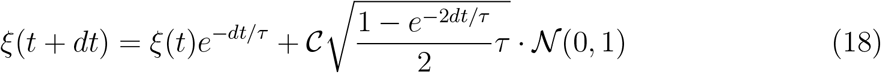

with *τ* being the correlation time and 𝒩(0, 1) a random normal distribution with mean zero and unitary variance. Furthermore, note that the variance of the OU process is related to the amplitude 𝒞 and the correlation time *τ* as

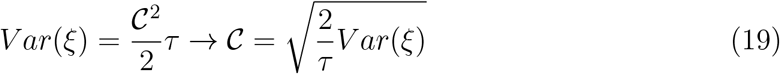

In order to analyze the effects of higher levels of correlations in the noise without changing the variance of the signal, we vary the amplitude accordingly by using the previous relation. The SVCA and PCA of white noise is reported in section 3, while the correlated noise is examined in Appendix A.

### 2.3. Experimental data sets

We have analyzed large-scale neural activity data sets recorded using calcium imaging methods in awake, spontaneously behaving mice. Specifically, we have considered two-photon microscopy data from the visual cortex (openly available at Stringer et al. (2018) and Syeda et al. (2023)) and light bead microscopy data from the dorsal cortex (available from Manley et al. (2024a)). In brief, the temporal fluorescence traces from calcium imaging represent continuous signals that serve as an indirect but widely used proxy for neuronal activity. These traces reflect a slow and noisy integration of action potential firing, capturing when and how strongly a neuron was active. Although calcium imaging methods have lower temporal resolution than direct electrophysiological recordings where one can accurately resolve the single neuron spikes, they allow for the simultaneous recording of hundreds to thousands of neurons. This enables robust estimation of correlations between the neurons’ activity patterns at the population level and allow for the detailed scaling analysis with the number of neurons reported in the following. Further detailed descriptions of the imaging methods can be found in the respective references. In Table 1 we detail the specific data sets used in our analysis, in particular the employed data sets will be indicated with the following acronyms : STk for Stringer et al. (2019b), SYk for Syeda et al. (2024) and MAk for Manley et al. (2024b) with *k* = 1, 2. In practice, in the paper are reported the analysis concerning the data sets numbered as #1, while the others numbered as #2 have been employed only to verify the consistency of the found results.

**Table 1:** Experimental data sets employed in our analysis. For each data set are specified the bibliographic and the specific data set reference, the total number of neurons, the time resolution at which the activity of the neurons are saved and the total number of datapoints, together with an acronym for each data set.

| reference | data set | total number<br>of neurons | total number<br>of datapoints | time<br>resolution | acronym |
| --- | --- | --- | --- | --- | --- |
| Stringer et al. (2019b) | spont M161025 MP030 20170623 | 14752 | 24354 | 0.4 s | ST1 |
| Stringer et al. (2019b) | spont M161025 MP030 20170616 | 15179 | 18136 | 0.4 s | ST2 |
| Syeda et al. (2024) | spont TX104 2022 10 05 2 | 60649 | 25193 | 0.3 s | SY1 |
| Syeda et al. (2024) | spont TX103 2022 10 05 2 | 84310 | 25583 | 0.3 s | SY2 |
| Manley et al. (2024b) | 20210310 left hemisphere FOV 289mW<br>50 550 $\mu\text{m}$ depth 30 min no stim | 315363 | 8640 | 0.2 s | MA1 |
| Manley et al. (2024b) | 20210310 right hemisphere 289mW<br>50 550 $\mu\text{m}$ depth 60 min no stim | 204798 | 17280 | 0.2 s | MA2 |

In practice and as described in the previous subsection 2.1.2, to perform SVCA the neurons are divided in two groups that we considered of the same size, and also the corresponding time samples are also split in half to perform training and test of the same duration. The quantities appearing in our scaling analysis will always refer to half of the total number of employed neurons and of the total duration, therefore these numbers will be always smaller than those reported in the Table 1. Furthermore, as it will become clear in the following, for the validation of our hypothesis we do not need to use the total amount of recorded neurons from each experiment, this is particularly true for the data reported in (Manley et al., 2024b).

The subsampling from the total number of neurons and the split of the population and timepoints in two sets for the implementation of the SVCA was carried out following the same procedures as described in the original papers. In particular, for the data sets of Stringer et al. (2019b) and Syeda et al. (2024), first the spatial XY plane of the recording is divided into *n* ≃ 20 non-overlapping strips of 60 *µm* each. Then, the population is split in two by assigning the neurons in the even strips into one group and the neurons in the odd strips to the other. In this way, there are no pairs of neurons in the two sets with the same spatial position but corresponding to different depths. This avoids a potential confound of two neurons predicting one another from its own out-of focus fluorescence, as explained in (Stringer et al., 2019b). Afterwards, the subsampling is done by randomly selecting *N* neurons from each subpopulation. On the other side, timepoints are separated by chunking the data into 72 seconds intervals and assigned alternatively to the training or test sets. For both these data sets we analyzed all the available neurons and timepoints.

For the data sets MA1 and MA2, the subsampling procedure was carried out using their own software provided alongside with the data in https://github.com/vazirilab/scaling_analysis, up to a maximum number of 65536 neurons. Their procedure consisted first in fixing a population size of 2 · *N* and subsampling this quantity from the full population randomly across the full field of view. Then, the population is split into the two sets by dividing the imaging field of view laterally into squares of 250 *µm* and grouping non adjacent squares in a checkerboard pattern. The aim of this strategy is the same as before, to prevent axial crosstalk between sets due to out-of-focus fluorescence (Manley et al., 2024b). The split of the timepoints was done similarly as for the other two data sets.

## 3. Results

Firstly, in order to characterize the SVC approach we will examine in subsection 3.1 synthetic noisy data in a high dimensional space generated by lifting the 3 dimensional dynamics of the Lorenz model in a *N* dimensional space in presence of additive noise. Moreover, we will investigate the ability and the drawbacks of the SVCA, as compared to PCA, to recover the reliable responses induced by the common underlying Lorenz dynamics.

In the following subsection 3.2 we will identify and clarify the main common features of the reliable and total SVC spectra for the experimental data sets described in subsection 2.3, which correspond to spontaneous neural activity in the visual and dorsal cortex in awake head fixed mice. Then we will focus on the dependence of the reliable and total variances {*S*_*k*_} and {*S*_*k,tot*_} on the number *N* of considered neurons and we will determine the asymptotic profile for the fraction of reliable variance 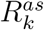 contained in the *k*-th SVC component. Finally, we will evaluate numerically and analytically the dimensionality of the reliable responses by employing different SVC-based approaches. For comparison we will report also the estimation of the PC-based embedding dimensions for the same data sets.

### 3.1. SVCA vs PCA for synthetic noisy data

In this subsection we will clarify the potentialities and limitations of the SVCA in recovering the dimensionality of a system subject to noise. As a peculiar example we will consider the well known Lorenz model (described in subsection 2.2) in presence of Gaussian white noise and we will compare the results of the SVCA with that obtained with a standard PC-based approach.

First, we consider the 3 dimensional Lorenz dynamics lifted onto a higher *N*-dimensional space via a linear transformation, as explained in subsection 2.2. We fix initially *N* = 500. Then, we add to this redundant high-dimensional system either uncorrelated white noise for different noise-to-signal ratios *ε*, as defined in Eq. (17), or correlated Ornstein-Uhlenbeck noise for different correlation times (this analysis is reported in Appendix A).

Let us start from the PCA. As shown in Fig. 1 (a) the value of the three first eigen-values is definitely larger than that of all the others for any amplitude *ε*, however their values decrease for increasing *ε* as expected. The first three directions are clearly associated to the Lorenz dynamics that in this case is the common underlying signal for all the *N* variables. In absence of noise we expect that the sum of the first three variances *λ*_*k*_ will correspond to the number *N* of variables, for increasing noise amplitude other directions with *k >* 3 will contribute more and more to the total variance. In particular, by assuming that the noise is affecting all the other directions in an equal manner, we expect that

**Figure 1:**
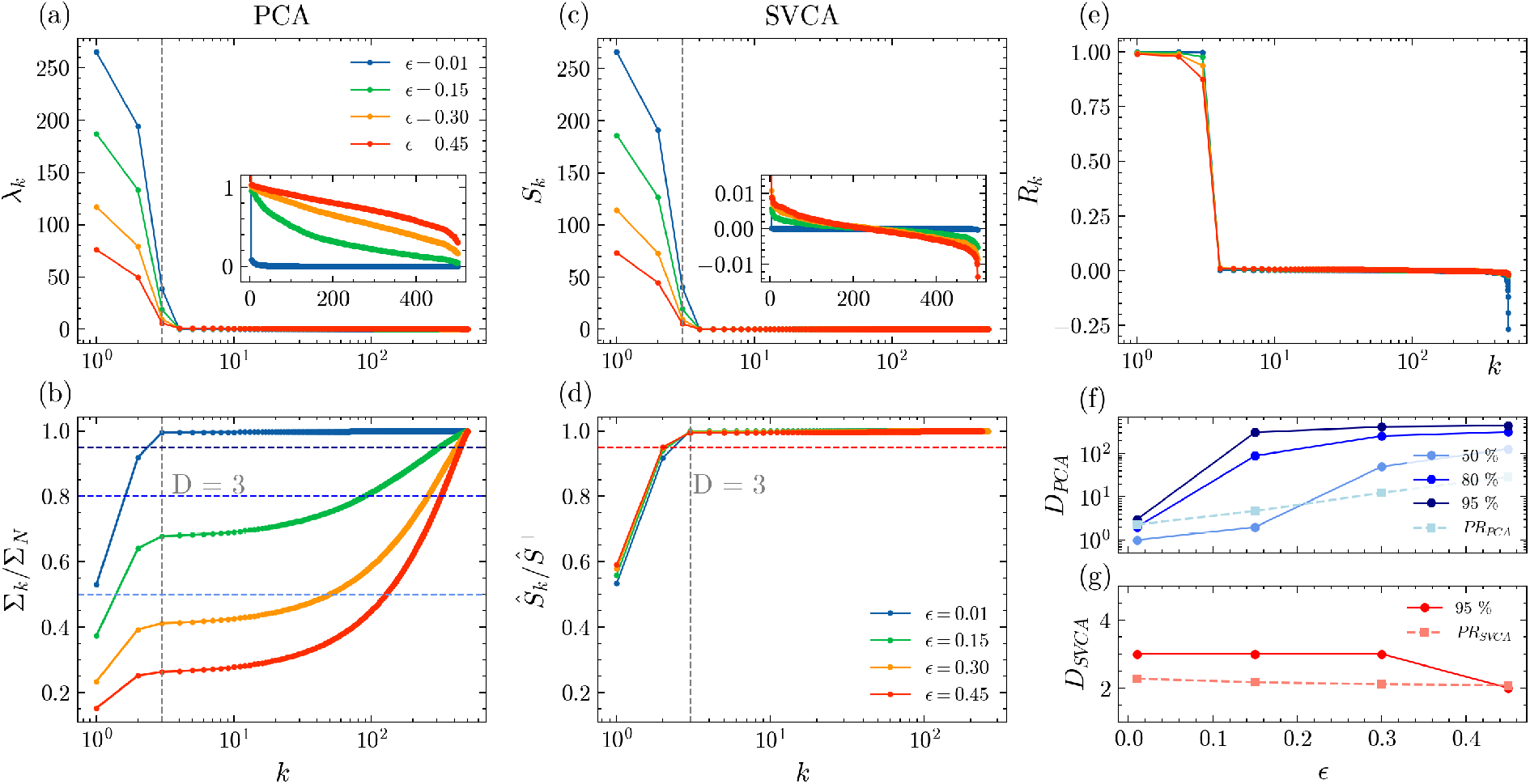
Lorenz system lifted onto a *N* = 500 dimensional space subject to additive white noise for different noise-to-signal amplitude ratios *ε*: namely *ε* = 0.01 (blue symbols), *ε* = 0.15 (green symbols), *ε* = 0.30 (orange symbols) and *ε* = 0.45 (red symbols). (a) Spectra of the PC eigenvalues *λ*_*k*_ versus their index *k*. An enlargement of the spectra for *k >* 3 is shown in the inset. (b) Normalized cumulative sum (variance) Σ_*k*_*/*Σ_*N*_ of the PC spectra. (c) Reliable SVC spectra *S*_*k*_. In the inset an enlargement of the spectra for *k >* 3 is shown. (d) Normalized cumulative sum *Ŝ*_*k*_ */Ŝ*^+^ of the reliable SVC spectra, please notice that the total sum *Ŝ*^+^ is limited to the positive *S*_*k*_. The vertical dashed line in (a-d) denotes the value of the dimension of the phase space of the Lorenz model *D* = 3 (e) Fraction of reliable variance contained in the *k*-th SVC. (f) System dimensionality *D*_PCA_(*p*) estimated by employing PC-variances as in Eq. (13) for *p* = 50% (medium blue), 80% (blue) and 95% (dark blue) versus *ε*. The colors refer to the horizontal dashed lines in (b). Also the participatio ratio *PR*_PCA_ (light blue squares) defined in Eq. (14) is reported. (g) Dimensionality *D*_SVCA_(*p*) (red symbols) associated to the reliable variance estimated by counting the SVC components for which *Ŝ*_*k*_ */Ŝ*^+^ *>* 0.95 (see panel (d)). In panel (g) it is also reported the *PR*_SVCA_ (light red square symbols) estimated by employing Eq. (14) limited to the the positive part of the reliable SVC spectrum *S*_*k*_, its value ranges from *PR*_SVCA_ ≃ 2.27 (*ε* = 0.01) to *PR*_SVCA_ ≃ 2.08 (*ε* = 0.45). The number of sampling time points has been fixed to *T* = 5 · 10^4^.

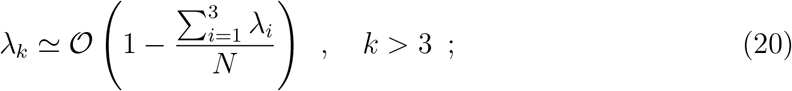

as we have numerically verified.

In order to estimate the corresponding embedding dimension *D*_PCA_(*p*) we consider the normalized cumulative spectra Σ_*k*_*/*Σ_*N*_ reported in Fig. 1 (b) and we fix three different values of *p*, namely *p* = 50%, 80% and 95%. As shown in Fig. 1 (f), *D*_PCA_(*p*) grows with *p* and approaches *N* for sufficiently large *ε* as expected for a system where the dynamics are dominated by noise. A similar increase is displayed by *PR*_PCA_. In particular, we observe that *PR*_PCA_ corresponds at low (high) noise to the value of *D*_PCA_(*p*) for *p* ≃ 60% (*p* ≃ 40%). Therefore the correspondence of *PR*_PCA_ with *D*_PCA_(*p*) occurs for values of *p* definitely smaller than those predicted by Gao et al. (2017) for any value of the noise amplitude. Furthermore, quite unexpectedly, the participation ratio for small noise *ε* = 0.01 gives a value of *PR*_PCA_ = 2.28, that is a good estimate of the intrinsic (correlation) dimension of the Lorenz attractor for these parameter values, namely *D*_*f*_ ≃ 2.05.

As shown in Fig. 1 (c), the SVC spectra {*S*_*k*_} are quite similar to {*λ*_*k*_}. The main difference is associated to the tail of the spectra *k >* 3, that is the most influenced by noise and where the values of *S*_*k*_ can even become negative, as shown in the inset of the figure. This is due to the fact that the singular basis employed to perform the singular value decomposition is estimated during the training period and therefore does not coincide with the basis of the cross-variance matrix obtained in the test period. For what concerns the cumulative spectra up to dimension *k, Ŝ*_*k*_, we observe in Fig. 1 (d) that they reach almost 100% of the cumulative sum *Ŝ*^+^ already at *k* = 3, and that the dependence on the noise amplitude is indeed almost negligible.

This is even more evident from Fig. 1 (e), where the fraction of reliable variance *R*_*k*_ corresponding to the *k*-th SVC dimension is reported. We observe that *R*_1_ ≃ *R*_2_ ≃ 1 for all the considered noise amplitudes and only *R*_3_ is affected by the noise-to-signal ratio amplitude *ε*, but with *R*_3_ *>* 0.80 up to *ϵ* = 0.45. Therefore, if we set a threshold to *p* = 95%, the embedding dimensionality associated to the reliable variables remains always *D*_*SVCA*_(*p*) = 3 for almost all the considered values of *ε*. It only drops to 2 for the largest noise amplitude, as shown in Fig. 1 (f), therefore in this case only the first two dimensions contain pure reliable signal unaffected by noise.

We have also measured the embedding dimensionality by employing the participation ratio of the spectrum *S*_*k*_, limited to the positive values, in this case we obtain *PR*_*SVCA*_ ≃2.27 − 2.08 depending on the value of *ε* not far from the intrinsic dimension *D*_*f*_ of the Lorenz attractor in absence of noise.

In order to understand the influence of noise on the SVC spectra and their dependence on the number of time points *T* and system size *N*, we have estimated the spectra for purely white noise signals. As shown in Fig. 2 (a), the reliable SVC spectra are symmetric around zero (the first *N/*2 values of *S*_*k*_ are positive and the remaining *N/*2 negative). The total SVC spectra (reported in the inset) are strongly fluctuating around zero.

**Figure 2:**
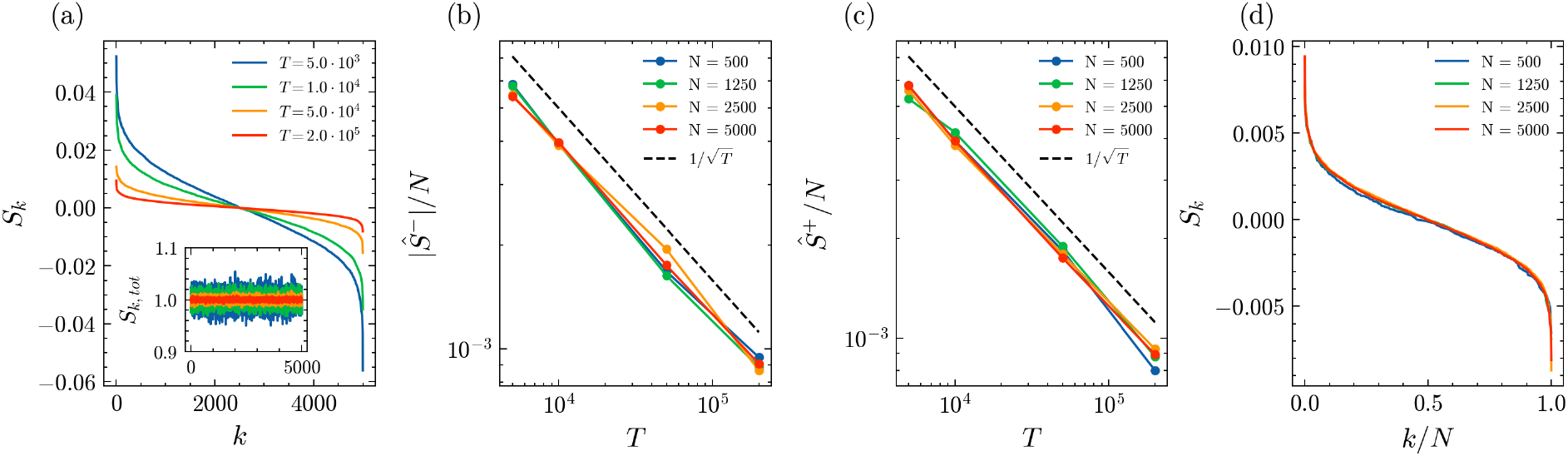
Pure white noise characterized in terms of the SVCA with standard deviation 𝒞= 5 corresponding to *ε* = 0.1 for the data reported in Fig. 1. (a) Reliable SVC spectra *S*_*k*_ measured for different number *T* of sampling data points for fixed system size *N* = 5000. In the inset are reported also the corresponding total SVC spectra *S*_*k,tot*_. The sum of the negative (positive) RVs |*Ŝ*^−^|*/N* (*Ŝ*^+^*/N*) is reported in panel (b) (panel (c)) rescaled with the system size *N* versus *T* for different system sizes. The dashed line denotes a decay 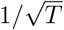. (d) Reliable SVC spectra *S*_*k*_ versus *k/N* for different system sizes *N* ranging from 500 to 5000 for a number of timepoints *T* = 2 · 10^5^.

Furthermore, by increasing *T* we observe that the values of the reliable and total SVC spectra shrink suggesting that *S*_*k*_ (*S*_*k,tot*_) will tend to zero (one) for *T* → ∞. The convergence of *S*_*k,tot*_ to one for sufficiently large *T* is expected since the total spectrum is associated to the covariance matrices of the two considered subpopulations and therefore it is akin to the PC spectrum, that for pure noise should be constant and exactly one along all directions. On the other hand, the values of the RVs *S*_*k*_ are associated to a cross-covariance matrix whose elements for noise signals are expected to vanish as 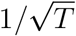 due to the central limit theorem. This latter result is confirmed by the data reported in Figs. 2 (b-c), where the sum of the positive (negative) part of the reliable spectra *Ŝ*^+^ (*Ŝ*^−^) is reported for different system sizes versus *T*. The simulations have been performed by considering an equal number of timepoints *T* in the training and test periods. However, to single out the relevance for the spectra convergence of either the training or the test period, we have analyzed training and test phases of different durations. The main results are that whenever the training period *T*_*train*_ is sufficiently long, in this case *T*_*train*_ ≥ 5000, its value does not influence the spectra and the decay of *Ŝ*^+^ and *Ŝ*^−^ with the test period duration *T*_*test*_ as 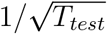 is still observable. These results remain true for *Ŝ*^−^ obtained for the lifted Lorenz model in presence of noise as discussed in the following.

Apart the vanishing of the reliable spectra as 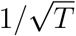, we observe that *Ŝ*^+^ (*Ŝ*^−^) grows linearly with *N*. This aspect can be explained by observing that for a fixed number *T* of sampling points the spectra *S*_*k*_ perfectly overlap for different system sizes if they are plotted as a function of *k/N*, as shown in Fig. 2 (d). This suggests that the spectra can take an universal form of the type *S*_*k*_ = *S*(*k/N*), therefore the cumulative sum of the positive (negative) eigenvalues can be rewritten as

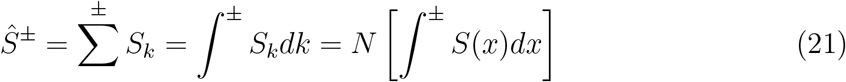

where *x* = *k/N* and the sums (integrals) are restricted to positive (negative) eigenvalues. Since the quantity appearing within the square parentheses on the rhs of Eq. (21) is independent of *N*, one expects that *Ŝ*^*±*^ will be extensive quantities, i.e. they should grow proportionally to *N*, and this explains the results reported in Figs 2 (b-c).

Let us now reconsider the noisy Lorenz data and in particular the dependence of the SVCA on the dimensionality *N* of the space where the Lorenz dynamics is lifted. As shown in Fig. 3 (a), in this case we observe that the sum of positive and negative values of the reliable SVC spectra *Ŝ*^+^ and *Ŝ*^−^ both grow proportionally to *N*. However, while *Ŝ*^−^ decreases as 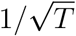 as in the pure noise case, *Ŝ*^+^ seems quite insensitive to the number of time-points and remains constant for increasing *T*. At the same time, for increasing system size there is a clear increase of the fraction of reliable variance *R*_*k*_ contained in the first three SVC components, *R*_*k*_ approaches one for *k* = 1, 2, 3 already for *N >* 500 (see Fig. 3 (b)).

**Figure 3:**
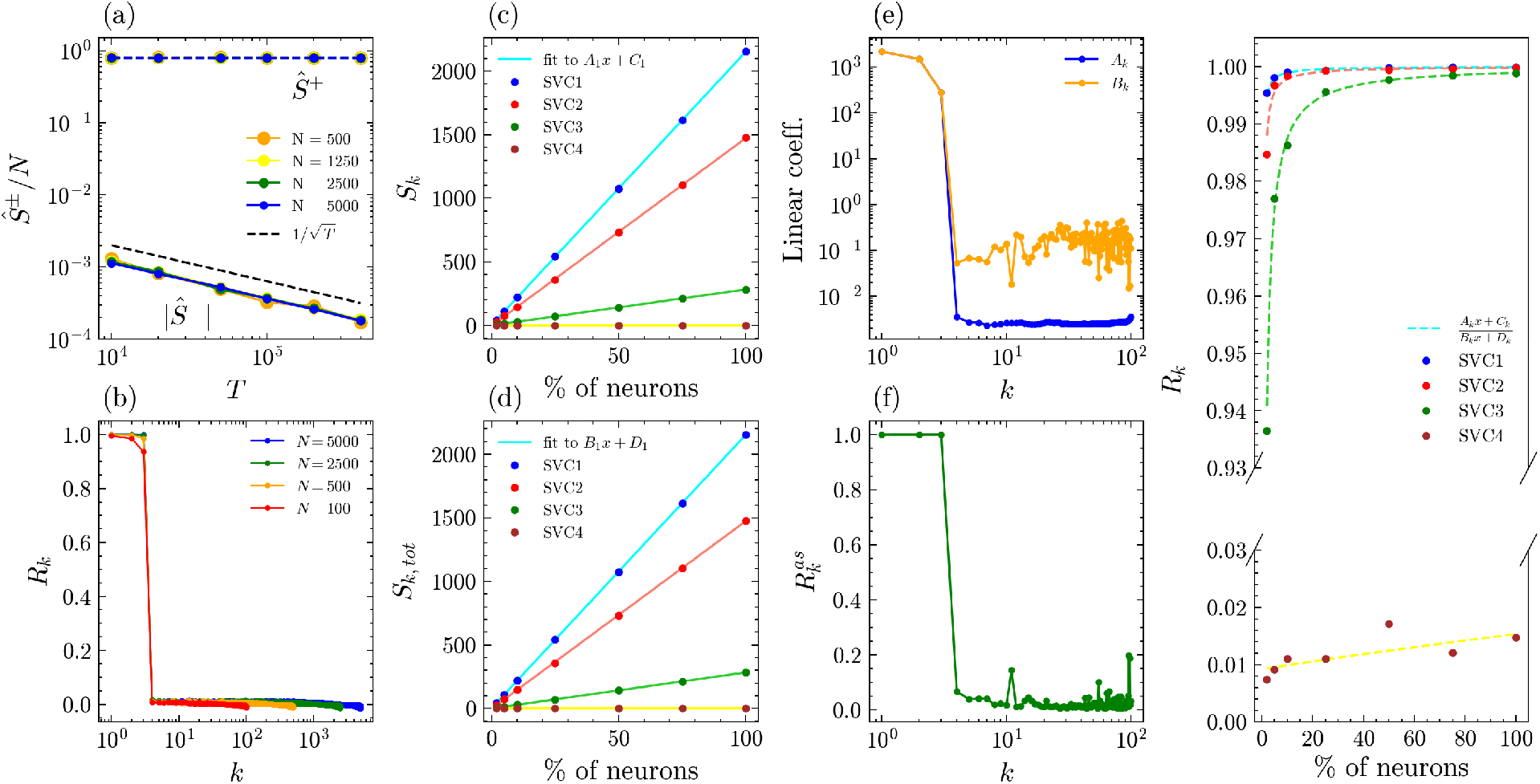
SVCA analysis of the Lorenz system lifted in an *N* dimensional space subject to white noise with noise-to-signal ratio *ε* = 0.1. (a) The sum of the negative (positive) SVC eigenvalues |*Ŝ*^−^| */N* (*Ŝ*^+^*/N*) rescaled with the system size *N* versus *T* for different system sizes. (b) Fraction of reliable variance contained in the *k*-th SVC for different system sizes and *T* = 5 · 10^4^. The first 4 SVCs *S*_*k*_ (c) and *S*_*k,tot*_ (d) are reported versus the % of considered neurons with respect to the maximal system size *N*_*m*_ = 5000 together with the linear fits *A*_*k*_*x* + *C*_*k*_ and *B*_*k*_*x* + *D*_*k*_. (e) Linear coefficients *A*_*k*_ and *B*_*k*_ versus *k* as obtained by fitting the data as a function of *N*. (f) Asymptotic values of the fraction of reliable variance 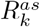 versus *k*. (g) *R*_*k*_ versus % of neurons for the first 4 SVCs.

These behaviours could be understood by analyzing the dependence on the system size of the first SVC eigenvalues *S*_*k*_ and *S*_*k,tot*_, these are reported in Fig. 3 (c) and (d) for different sizes *N*, expressed as a % of a maximal size fixed in this case to *N*_*m*_ = 5000. For *k* = 1, 2, 3 we observe a linear increase with the number of elements *N* = %*N*_*m*_, that can be expressed as

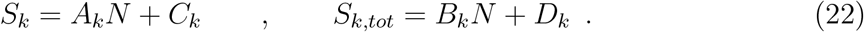

The corresponding linear coefficients *A*_*k*_ and *B*_*k*_ obtained by the fits to the data are of the order of 10^2^ − 10^3^ for the first three SVCs and then drop to values of order 𝒪 (10^−3^) (𝒪 (10^−1^)) for *A*_*k*_ (*B*_*k*_) for *k >* 3, see Fig. 3 (e). In particular, the large fluctuations in the values of *B*_*k*_ observable at *k >* 10 are due to the fact that the linear scaling with *N* for this quantity becomes more and more questionable for large *k*.

The linear increase with *N* of the values of the first three variances *S*_*k*_ and *S*_*k,tot*_ can be explained by the fact that they encode for the three variables of the Lorenz model, that represent the collective global variables controlling the whole dynamics. Furthermore, the fact that the values of *Ŝ*^+^*/N* are independent by *N*, as shown in Fig. 3 (a), is due to the fact that the first three coefficients *A*_*k*_ are definitely larger than the others. Therefore the sum *Ŝ*^+^ is dominated by few elements growing with *N*, while the contribution of the rest of the SVC dimensions to this sum is negligible. The scaling of *Ŝ*^−^ with *N* has instead a different origin due to the extensive property of the negative part of the spectrum (see Eq. (21)) analogously to what we found for pure noise.

If now we consider that the fraction of reliable variance, this can be written as

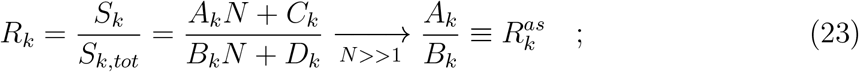

where the ratio 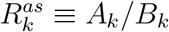 represents the asymptotic value of *R*_*k*_ that can be achieved for sufficiently large system sizes. This ratio is shown in Fig. 3 (f) and reveals that for sufficiently large *N* the first 3 SVC dimensions encode only the reliable part of the variance 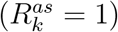. For larger *k* the ratio 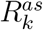 drops to very small values indicating these further components are essentially associated to noise.

These results are confirmed by considering directly *R*_*k*_ versus *N* (shown in Fig. 3 (g)) for the first 4 SVC dimensions. While for *k* = 1, 2, 3 the fraction *R*_*k*_ approaches 100 % for *N* = 5000 (the maximal size here considered), at this size *R*_4_ barely reaches a value slightly larger than 1 % and it shows a tendency to saturate.

### 3.2. SVC vs PC-characterization of spontaneous neural activity

As already mentioned, SVCA has been introduced in (Stringer et al., 2019b) to characterize spontaneous activity in mouse visual cortex, and more recently SVCA has been employed in (Manley et al., 2024b) to show an unbounded growth of reliable dimensionality with the number of neurons recorded simultaneously from the whole mouse cortex.

We will re-examine critically in terms of SVCA and PCA some of the available data sets provided in (Stringer et al., 2019b; Syeda et al., 2024; Manley et al., 2024b): namely, we will consider ST1, SY1, and MA1, as defined in Table 1.

#### 3.2.1. Dependence of the SVC-results on the number N of neurons

Let us first perform the SVCA for different number *N* of neurons, obtained by resampling the considered data sets as explained in subsection 2.3, in order to identify characteristic aspects. As evident from Figs. 4 and 5 such data sets share the following common features :

**Figure 4:**
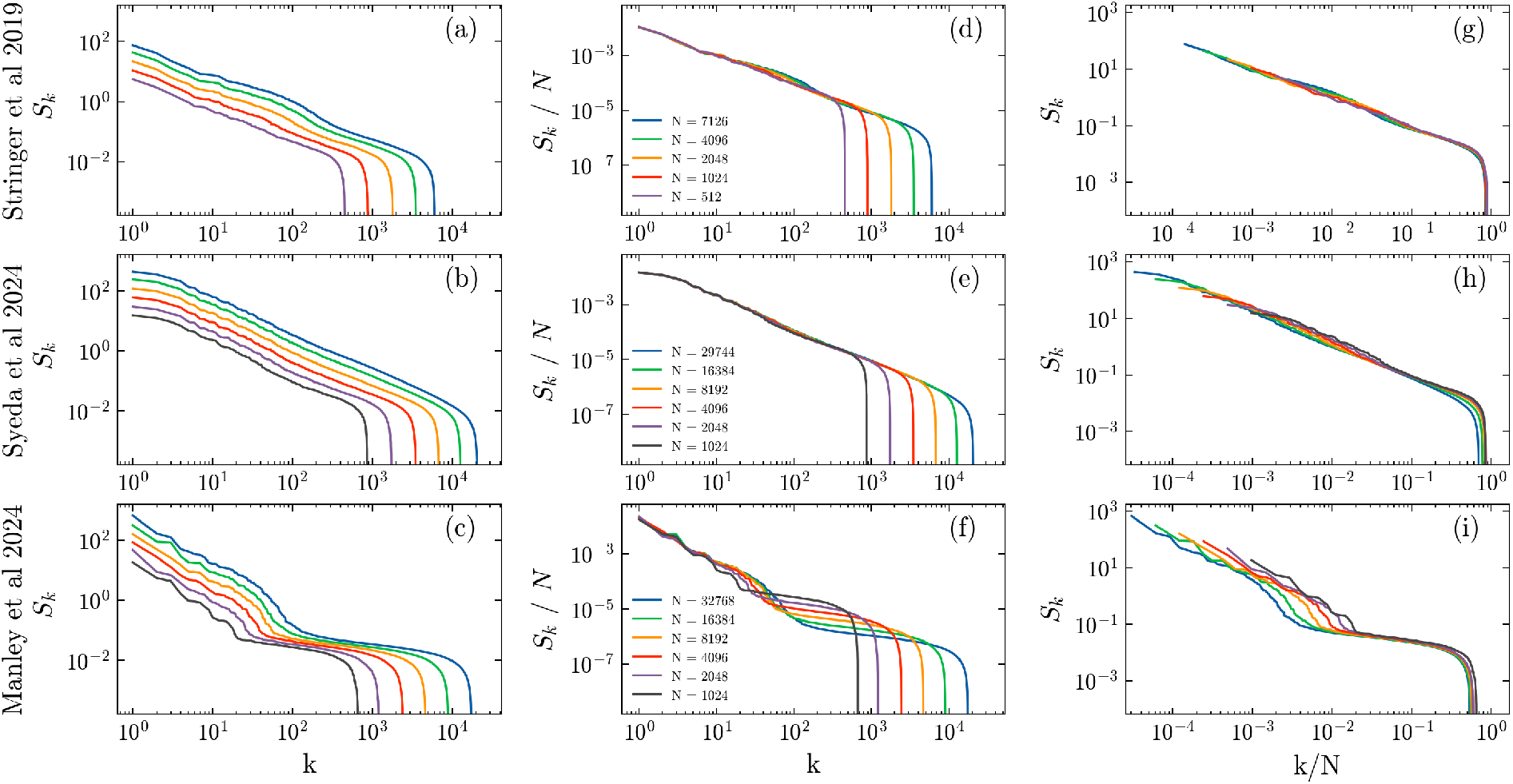
Experimental data for the RVs {*S*_*k*_} corresponding to data sets ST1 (upper row), SY1 (middle row), and MA1 (lower row) reported for different numbers *N* of examined neurons, indicated in the legends displayed in the second column. The RVs *S*_*k*_ versus *k* are shown in the first column, *S*_*k*_*/N* versus *k* are reported in the second one and *S*_*k*_ versus *k/N* in the third one.

**Figure 5:**
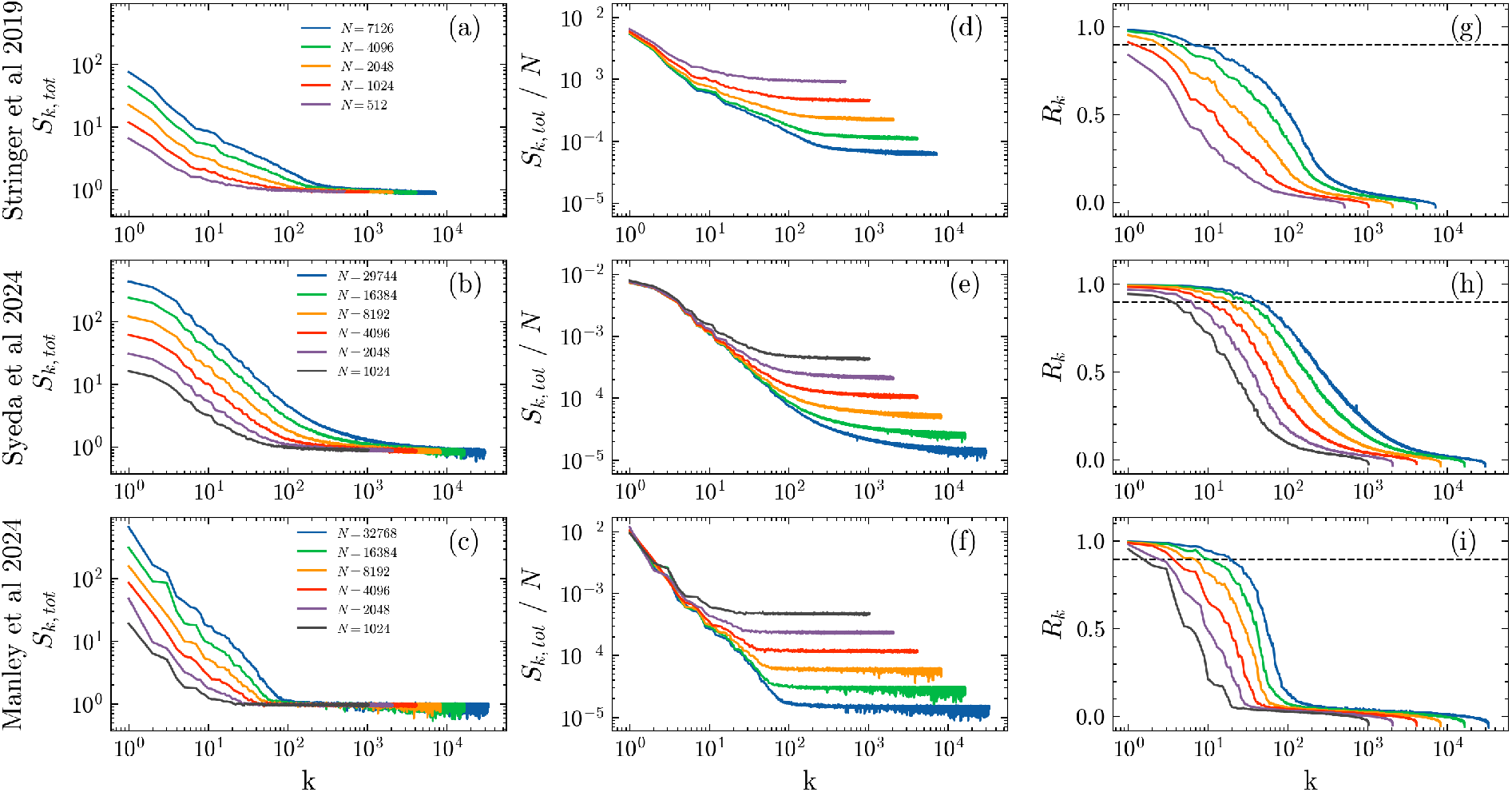
Experimental data for the TVs {*S*_*k,tot*_} and the fractions of reliable variance {*R*_*k*_} corresponding to data sets ST1 (upper row), SY1 (middle row), and MA1 (lower row) reported for different numbers *N* of examined neurons, indicated in the legends displayed in the second column. The first column displays the TV *S*_*k,tot*_ versus *k*, the second one reports *S*_*k,tot*_*/N* versus *k* and the third one *R*_*k*_ versus *k*, with the black dashed line denoting *R*_*k*_ = 0.9.

i. the reliable variances *S*_*k*_ display a power law decay *k*^−*α*^ up to some critical SVC dimension *k*_*c*_ = *k*_*c*_(*N*), which grows with *N*, as evident from the data reported in the first column of Fig. 4;
ii. the total variances *S*_*k,tot*_ saturate to an almost constant value 𝒪 (1) above some 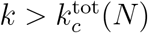, which grows with *N*, as shown in the first column of Fig. 5.
iii. the variances *S*_*k*_ and *S*_*k,tot*_ associated to the first SVC components grow proportionally to the number of considered neurons *N* (as shown in the second column of Figs. 4 and 5);
iv. the RVs *S*_*k*_ associated with the last SVC components essentially collapse one on the other when reported as a function of *k/N* ^1^, namely *S*_*k*_ = *S*_*k*_(*k/N*) for sufficiently large *k > k*_*c*_(*N*), as clear from the data displayed in the third column of Fig. 4;
v. The number of SVCs with a fraction of reliable variance *R*_*k*_ larger than a certain arbitrary threshold Θ increases with the number of neurons *N*, as shown in the third column of Fig. 5

For what concerns the differences among the data sets, apart the power-law exponent *α* that depends on the considered set, a peculiar difference is that for MA1 *S*_*k*_ for *k > k*_*c*_(*N*) display a clear plateau before dropping to extremely low and even negative values, a plateau not present in the other 2 data sets.

In the following we will examine in details the common aspects i)-v) delineate above.

i. -iv) The power-law decay of the reliable variances have been reported in several different neural experiments concerning the cortex activity of awake mice under stimulation (Stringer et al., 2019a) and during spontaneous activity (Stringer et al., 2019b; Stringer and Pachitariu, 2024; Manley et al., 2024b). The value of the power law exponent *α* is believed to be related to the dimensionality of the space of the inputs for stimulus-evoked activity (Stringer et al., 2019a) and it has been found to be *α* ≃ 1 in movement-driven activity (Stringer et al., 2019b). We measured the power-law decay exponents for the considered data sets, which are always relative to spontaneous activity, and the corresponding values are reported in Table 2. The variability in the measured exponent for each data set emerges by considering the subsamplings of different size *N* reported in Fig. 4. We will assume that the only significant part of the reliable spectra is the one showing a power law decay. The remaining part of the spectra at *k > k*_*c*_(*N*) is essentially due to unreliable and/or noisy activity. This claim is supported by the point iv) in the above list, since the scaling of the reliable spectra observable at *k > k*_*c*_(*N*) is the same as the one observed for pure noise, reported in Fig. 2 (b).
ii. The saturation of *S*_*k,tot*_ to an almost constant value of order 𝒪 (1) is due to noise present in the neural signals, as already emerged in the discussion of pure noisy data, see the inset in Fig. 2 (a). In particular, we expect that

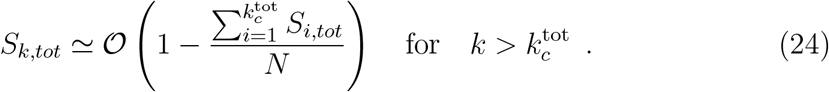 Furthermore, the fact that 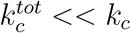 clearly indicates the effectiveness of the SVCA approach, which is able to extract information about reliable components even when the TVs are saturated to an almost constant value with apparently no information content.
iii. The direct proportionality of the values of *S*_*k*_, for *k* ≤ *k*_*c*_(*N*), and of the *S*_*k,tot*_, for 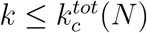, with the number *N* of neurons indicates that the corresponding SVCs encode for *collective* unobservable state variables influencing the dynamics of all the considered neurons, as it occurs for the first 3 SVCs for the lifted Lorenz model, see Fig. 3 (c-d). This is a further indication that the first SVC dimensions are not dominated by noise, but indeed they represent reliable states of the mouse.

v) As we will show in the following subsection 3.2.2 the fraction of reliable variances tend to an asymptotic profile 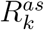 (23), therefore the number of SVCs for which *R*_*k*_ *>* Θ cannot grow in an indefinite manner with *N* and it should first or late saturate.

**Table 2:**
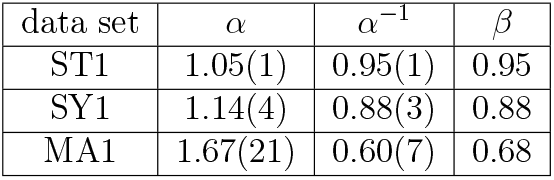
For the considered experimental data sets, we report the power-law decay exponents *α* of the RV spectra with their inverse *α*^−1^ and the exponent *β* controlling the power-law growth of the maximal reliable SVC *k*_*c*_ = *k*_*c*_(*N*) with the number of neurons *N*.

| data set | $\alpha$ | $\alpha^{-1}$ | $\beta$ |
| --- | --- | --- | --- |
| ST1 | 1.05(1) | 0.95(1) | 0.95 |
| SY1 | 1.14(4) | 0.88(3) | 0.88 |
| MA1 | 1.67(21) | 0.60(7) | 0.68 |

#### 3.2.2. Asymptotic results (N → ∞)

As already mentioned and as shown in the second columns of Figs 4 and 5, the dependence of *S*_*k*_ (*S*_*k,tot*_) on the number *N* of neurons is definitely linear for at least several hundreds (tenths) of SVCs. This scaling is clearly shown in Figs. 6 (a-c) for the SY1 data set, but this behaviour is common to all the examined data sets. As done for the Lorenz model, we can fit *S*_*k*_ and *S*_*k,tot*_ with the expressions (22) and obtain the corresponding linear coefficients *A*_*k*_ and *B*_*k*_ (shown in Fig. 6 (d)), as well as the asymptotic values for the fraction of reliable variance 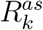 reported in Fig. 6 (e). This linearity holds for *k* even as large as 900 as we can see from Fig. 6 (c), where the fits reproduce well the scaling for each displayed SVC.

**Figure 6:**
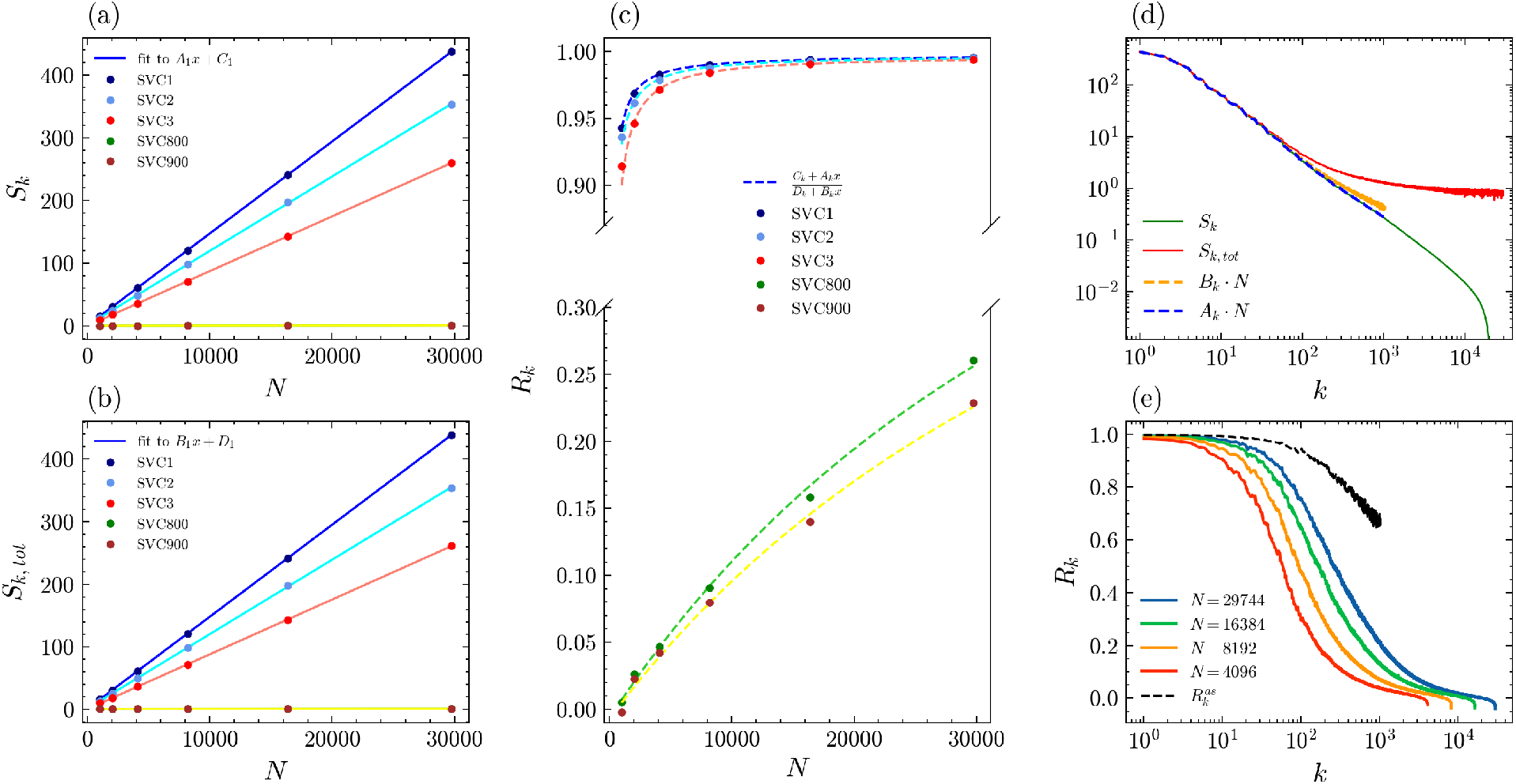
SVCA analysis of neuronal data set SY1. Reliable (a) and total (b) variances *S*_*k*_ and *S*_*k,tot*_ as a function of the number *N* of neurons for the first 3 SVCs and for higher SVCs. The solid lines correspond to linear fits to each SVC. (c) Fraction of reliable variance *R*_*k*_ against system size for the same SVCs as in (a-b). Dashed lines correspond to the ratio of the fittings reported (a,b). (d) *S*_*k*_ (green solid line) and *S*_*k,tot*_ (red solid line) together with *A*_*k*_ × *N* (blue dashed line) and *B*_*k*_ × *N* (dashed yellow line) versus *k* for *N* = 29744. (e) Fraction of reliable variance *R*_*k*_ versus *k* for different number of neurons and with the black dashed line denoting 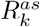.

The meaning of the asymptotic values 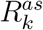 can be understood by comparing the spectra *S*_*k*_*/N* (*S*_*k,tot*_*/N*) obtained for some finite size *N* with the asymptotic values *A*_*k*_ (*B*_*k*_). Let us consider these quantities reported in Fig. 6 (d) for *N* = 29744 for the SY1 data set. First of all we notice that in the shown *k*-range, *S*_*k*_ (green solid line) is essentially coincident with *A*_*k*_ · *N* (blue dashed line), thus indicating that the intercept *C*_*k*_ obtained by the linear fit reported in (22) is negligible for this system size. On the other side, while *S*_*k,tot*_ (red solid line) is saturating to a constant value 𝒪 (1) at a sufficiently large *k*, the quantity *B*_*k*_ · *N* is essentially coincident with *S*_*k*_ up to *k* ≃ 200, and then assumes values slightly larger than the RVs. Therefore, the extrapolation to the asymptotic values as *N*→ ∞ *removes* some of the the unreliable components present in *S*_*k,tot*_, presumably those due to finite size fluctuations, while this removal has been already effectively achieved in *S*_*k*_ thanks to the SVC procedure. This explains why the asymptotic values of 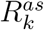 are definitely larger that the actual values *R*_*k*_ even for the largest system size here considered (namely, *N* = 29744). Indeed, at *k* = 2048, corresponding to the largest SVC at which we could perform the extrapolation, we obtain 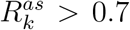 while *R*_*k*_ ≃ 0.1 (as shown in Fig. 6 (e)).

Unfortunately, we cannot perform reliable linear fitting (22) beyond *k* = 2048 in the present case. Since the saturation of *S*_*k,tot*_ for 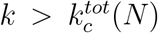 renders the linear scaling questionable and induces enormous fluctuations in the estimation of the parameter *B*_*k*_ and therefore also of 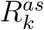. This can can be deduced from the increase in the dispersion of the orange curve in Fig.6 (d). Despite this limitation, the fact that the fraction of reliable variance 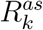 is not 100 % for all the SVCs even in the limit of infinite neurons indicates that the unreliable components cannot be completely removed in *S*_*k,tot*_ even in this limit.

#### 3.2.3. Reliable dimensionality of the neural data sets

Let us now examine how the dimensionality of the phase space associated to the reliable responses could be estimated by following different approaches.

A first approach, reported e.g. in Manley et al. (2024b), consists in identifying a critical value of the SVC dimension 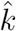 at which 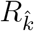 becomes smaller than a fixed threshold Θ and to find how 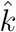 scales with the number of neurons *N*. In particular, Manley et al. (2024b) reported, a power-law growth of the neural dimensionality given by 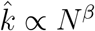 with an exponent *β* = 0.66(6), independently of the chosen threshold Θ.

We have performed the same analysis by setting Θ = 0.90 for SY1 and MA1 data sets. As shown in Fig. 7 (a-b), our analysis confirms a power-law growth of 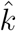 with *N* with exponents *β* ≃ 0.88 and *β* ≃ 0.60, at least for sufficiently low values of *N*. However, as evident from panel (a), for the SY1 data at *N* = 29744 we observe a deviation from this growth suggesting a saturation of the data. This is even more evident for the MA1 data shown in panel (b), where we have reported also the data taken from Fig. S3 (e) in (Manley et al., 2024b) (shown as green stars). Here, 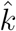 clearly bends from the power-law growth at *N* = 10^6^. In Fig. 7 (a-b) we report also, as horizontal dashed lines, the asymptotic values of 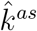 where 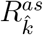 becomes smaller than 0.90, which correspond to 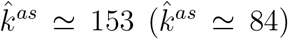 for SA1 (MA1) data set. These values are consistent with the displayed tendencies to a saturation of the values of 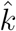 for growing *N*. From this analysis we conclude that an unbounded growth of the dimensionality with the neuron number (Manley et al., 2024b) is not plausible and a saturation should be expected.

**Figure 7:**
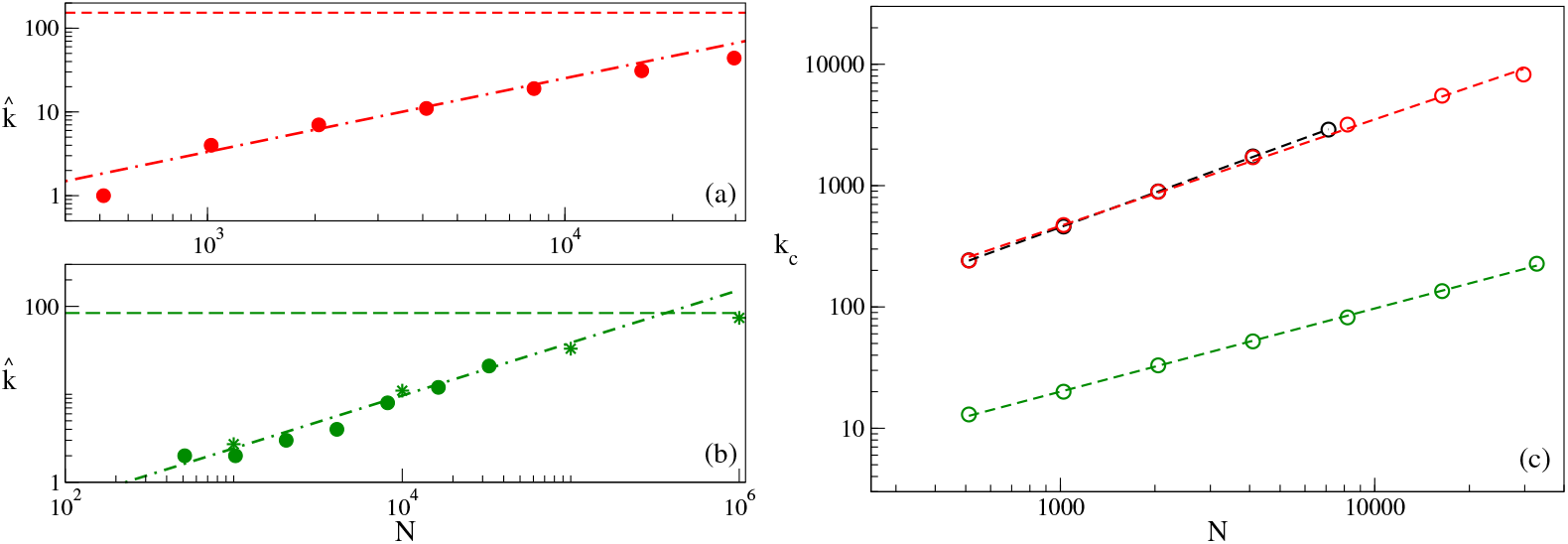
(a-b) Critical values 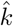 of the SVC at which 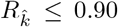. The results refer to SY1 (a) and MA1 (b): circles are our estimations, stars in (b) have been taken form Fig. S3 (e) in (Manley et al., 2024b). The dashed horizontal lines correspond to the asymptotic values of 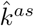 obtained by estimating the minimal SVC such that 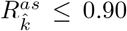. The dot-dashed lines refer to power-law scaling of the type 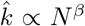 with *β* = 0.88 (a) and *β* = 0.60 (b). (c) *k*_*c*_ versus *N* for ST1 (black circles), SY1 (red circles) and MA1 (greeen circles). The definition of *k*_*c*_ is reported in the text. Dashed lines denote power law fitting to the data as *k*_*c*_ ∝ *N*^*β*^. The value of the estimated *β* exponents are 0.95 (ST1), 0.88 (SY1) and 0.68 (MA1).

In order to understand the origin of the exponent *β* controlling the initial power-law growth of 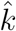 with *N*, we will now consider the dependence on the system size of *k*_*c*_(*N*), previously defined as the largest SVC for which a power-law decay is still observable in *S*_*k*_.

In practice, the value of *k*_*c*_(*N*) has been determined by fixing a quite small threshold Θ and then by estimating the largest *k* for which the RV *S*_*k*_ ≤ Θ. The thresholds have been chosen by analyzing for which value *S*_*k*_ bends from the power-law decay in Figs. 4(a-c). This resulted in threshold values of Θ = 0.02 for ST1 and SY1, while, due to the different profile of the *S*_*k*_, we set Θ = 0.06 for MA1. It is important to stress that for 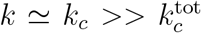, we have that *R*_*k*_ ≃ *S*_*k*_, since *S*_*k,tot*_ ≃ 1 for such SVCs. Therefore, we can safely affirm that (for sufficiently small Θ) our procedure is analogous to the one reported in (Stringer and Pachitariu, 2024) and (Manley et al., 2024b) to estimate the reliable dimensionality.

The scaling of *k*_*c*_ with *N* can be simply found by assuming that

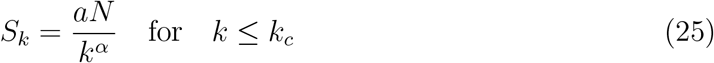

by following the results reported in points i) and iii) above. Therefore *k*_*c*_ should depend on the number of neurons as follows

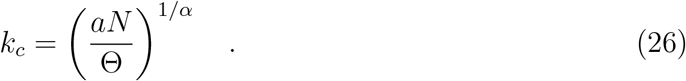

A numerical fit to the data with a power-law of the type *k*_*c*_ ∝ *N* ^*β*^ shown in Fig. 7 (c) gives estimations consistent with the forecasted scaling (26): namely, *β* ≃ 0.95 versus 1*/α* = 0.95(1) for ST1, *β* ≃ 0.88 versus 1*/α* = 0.88(3) for SY1, and *β* ≃ 0.68 versus 1*/α* = 0.60(7) for MA1 (these values have been also reported in Table 2). As evident from Fig. 7 (c), the results for the data sets ST1 and SY1 are almost identical up to *N* ≃ 16000, while the data for MA1 are definitely smaller, this is due to the differences in the power-law decay exponent *α* of *S*_*k*_. Furthermore, the power-law growth reported in (Manley et al., 2024b) with an exponent *β* = 0.66(6) is in perfect agreement with the estimate reported here for the MA1 data set, confirming the equivalence of our approach and that reported in (Manley et al., 2024b).

Even if the actual values of *k*_*c*_ clearly depend on the chosen threshold (26), the scaling law with the number of neurons will be not affected by the choice of it (as was also shown in (Manley et al., 2024b)). This is true at least as *k*_*c*_ are far from their corresponding saturation values, that we expect to be extremely large due to the small threshold values here considered. In summary, the observed scalings of 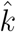 and *k*_*c*_ with *N* are simply due to the power-law decay of *S*_*k*_ (25).

The value of 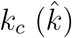 represents the number of SVC components that can be considered as reliable, since the corresponding *S*_*k*_ (*R*_*k*_) are above a certain threshold Θ for *k* ≤ *k*_*c*_ 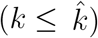. However, the estimation of the linear dimension for a given data set is usually related to the values taken by the variances along such components, the so-called embedding dimension in the definition reported in (Jazayeri and Ostojic, 2021). Components with vanishingly small variances contribute only marginally to the dimensionality of the linear embedding space.

As explained in subsection 2.1.3, the estimation of the embedding dimension is usually performed by employing two main methods: the standard one which gives the number of components *D*(*p*) needed to reconstruct a percentage *p* of the total variance, and the participation ratio *PR*, that is commonly used in physics to quantify *localization* (Bell and Dean, 1970), and does not require choosing an arbitrary threshold for the reconstructed variance.

We have numerically estimated *PR*_*SVCA*_ and *DP*_*SVCA*_(*p*) for the RVs spectra {*S*_*k*_}. In the following we limit to consider the SY1 and MA1 data sets, since the results for ST1 are quite similar to those of SY1, but limited to a smaller number of neurons. We report the outcome of this analysis as a function of the number of neurons in Figs. 8 (a-b) and (e-f). There are two peculiar differences among such data sets. Firstly, there is a clear dependence on *N* for SY1, while for MA1 the estimated dimensions are barely dependent on the number of neurons. A second aspect is that the dimensionality of the MA1 data is definitely smaller than those of the SY1 data. For example for SY1 data the *PR*_*SVCA*_ grows from 19 at *N* = 512 to 34 at *N* = 29744, while for MA1 data in a similar range the participation ratio takes values from 4 to 6. Similarly, we can observe that *D*_*SVCA*_(*p*) (for *p* = 70%) grows from 19 to 98 for SY1 data, while it takes value from 4 to 10 for the MA1 data in the range *N* ∈ [512, 32000]. Another interesting aspect to stress is that the values of *PR*_*SVCA*_ are comparable to those obtained with the *DP*_*SVCA*_(*p*) when *p* ≃ 60% for both SY1 and MA1 data (see Figs. 8 (a) and (e)) similarly to what observed for the noisy Lorenz data in subsection 3.1.

**Figure 8:**
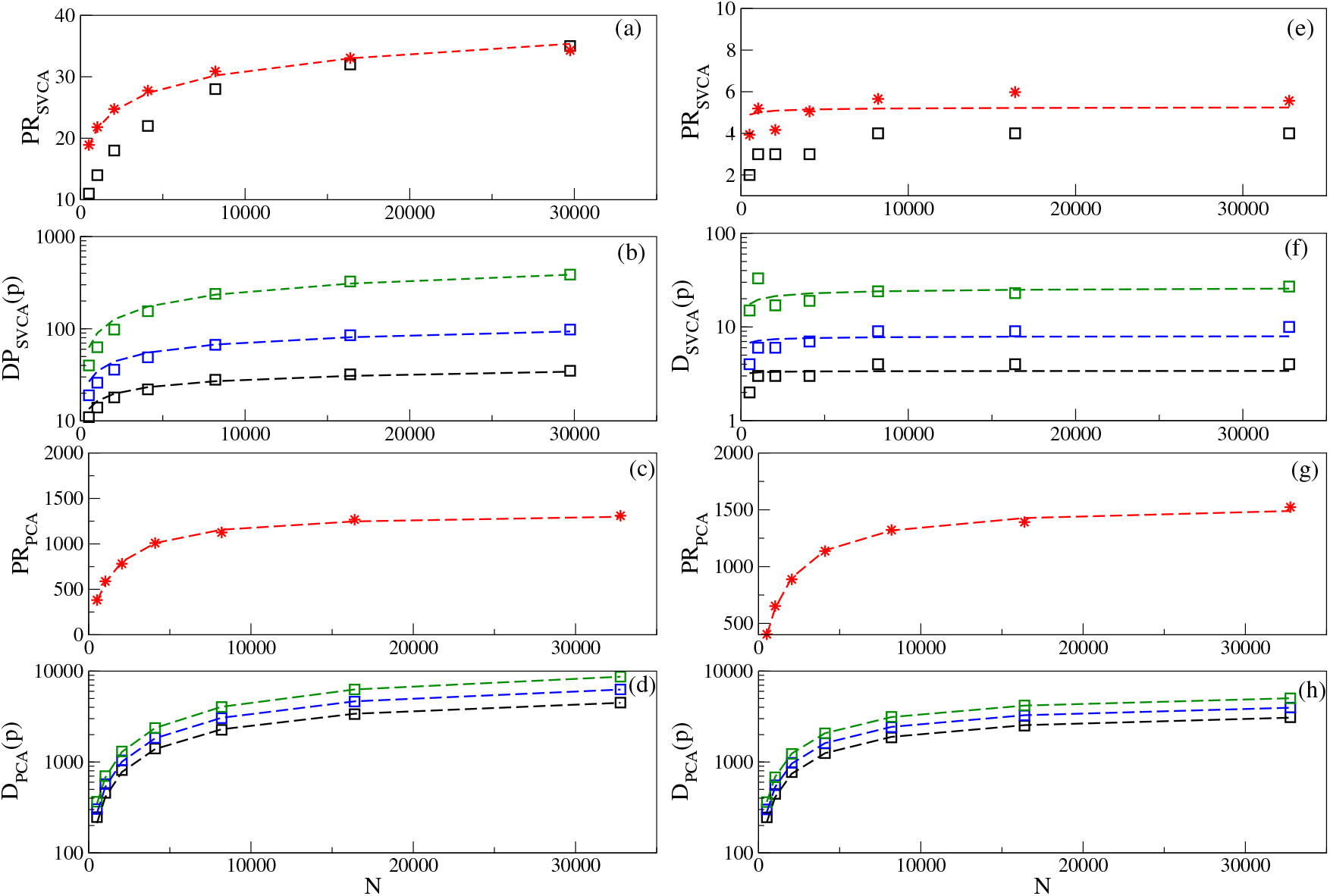
SVC and PC-based estimations of the data dimensionality: the first column (panels (a-d)) refer to SY1 data, the second column (panels (e-h)) to MA1 data. The *PR*_*SVCA*_ versus *N* (red stars) together with the fitting to the theoretical expression (27) (dashed red line) are reported in (a) and (e). In these panels are also reported the corresponding numerical estimate of the embedding dimension *D*_*SVCA*_(*p*) for *p* = 60% (black squares). The embedding dimensions *D*_*SVCA*_(*p*) (squares) versus the number of neurons *N* are shown together with the fitting to the theoretical expression (28) (dashed lines) in (b) and (f). The *PR*_*PCA*_ versus *N* (red stars) are reported in panel (c) and (g) with the fitting to the expression (30) (dashed red line). The *D*_*PCA*_(*p*) (squares) are reported with the fitting to the expression (30) (dashed lines) in (d) and (h). The colors in (b),(d),(f) and (h) denote the value of *p*: namely, *p* = 60% (black), 70% (blue) and 80% (green).

As we have shown, the actual value of the embedding dimension can depend on the method employed, and for *D*_*SVCA*_(*p*) also on the fraction *p* of the variances of the neural activity that we want to capture with the linear reconstruction. However, a fundamental question is if the dimension will grow unbounded with the number of neurons (Manley et al., 2024b) or if its value will saturate for sufficiently large *N*. To answer this question we have derived analytical expressions in Appendix B for the participation ratio and *D*(*p*) as a function of the number *N* of neurons. In particular, for spectra decaying as a power-law {*a/k*^*α*^} with *k* = 1, …, *N*, as the *S*_*k*_ spectra here considered (25), the expressions are the following ones

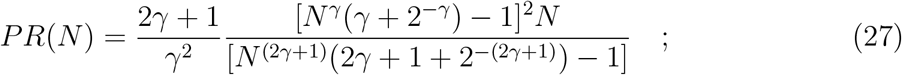

and

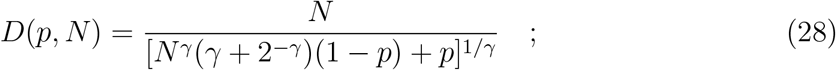

where *γ* = *α* − 1.

The comparison of these analytical expressions and the numerical data is reported in Figs. 8 (a) and (e) for the *PR*_*SVCA*_ and in Figs. 8 (b) and (f) for the *D*_*SVCA*_(*p*). In practice, we have fitted the numerical data with the expressions (27) and (28) in order to estimate *α*, the only free parameter appearing ins such equations. Then, we have verified the consistency of these fitted values *α*^*f*^ with the *α* exponents characterizing the power-law decay of *S*_*k*_ (see Table 2). For what concerns the participation ratios, the fitting to the data gives *α*^*f*^ ≃ 1.10 and *α*^*f*^ ≃ 1.49 for the SY1 and MA1 data, respectively, in agreement with the values reported in Table 2. For the *D*_*SVCA*_(*p*), we considered data for *p* = 60%, 70% and 80%. In this case, the estimated *α*^*f*^-values decrease with *p* and slightly overestimates the power-law decay exponents for the SY1 data. Indeed, we have obtained *α*^*f*^ = 1.18(1), while for MA1 the obtained values cover a wider but compatible range with the corresponding power-law decay exponents, namely *α*^*f*^ = 1.50(7).

Once verified that the above expressions (27) and (28) give reasonable estimates for the neural data, we can employ such expressions to verify if in the limit of an infinite number of neurons they will converge to a constant value or not. The corresponding asymptotic values obtained in the limit *N* → ∞ for *α >* 1 reads as

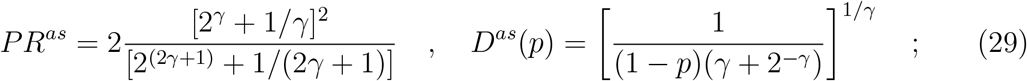

which depend only on the parameters *α* and *p*, as expected. The asymptotic values of the embedding dimensions obtained by employing the fitted parameters *α*^*f*^ are all finite since *α*^*f*^ *>* 1 in all the considered cases (see Table 3). More specifically for SY1 we obtain *PR*^*as*^ ≃ 89.6, while for MA1 *PR*^*as*^ ≃ 5.3, with the two quantities differing by more than one order of magnitude. On the other hand, the estimated asymptotical values of 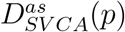 for SY1 (MA1) range from 89.2 (3.4) for *p* = 60% to 4187.3 (27.9) for *p* = 80%, confirming the fact that the estimated embedding dimensions for the SY1 data set are definitely larger than those for the MA1 data set at any value of *N*.

**Table 3:**
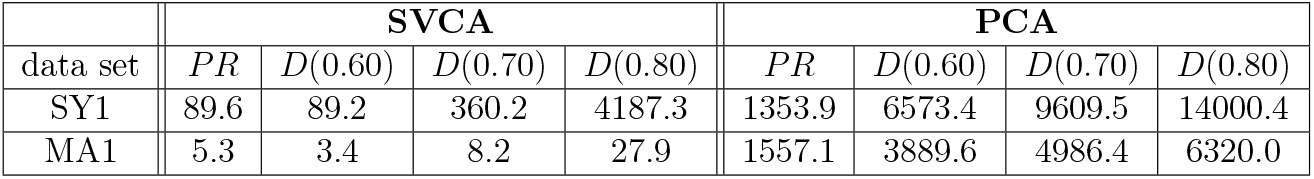
Asymptotic values for the embedding dimensions *PR*^*as*^ and *D*^*as*^(*p*) for the SVCA and the PCA for SY1 and MA1 data sets. For a better readability the superscript *as* is dropped in the table.

#### 3.2.4. PC-based embedding dimensionality of the neural data sets

Let us now consider the PC-based embedding dimensionality of the SY1 and MA1 data sets. Analogously to the SVCA, we have employed the *z*-scored neural activities to obtain PC-based dimensionality estimates via the participation ratio *PR*_*PCA*_ and *D*_*PCA*_(*p*) for various values of *p*. The numerical results for different number *N* of neurons are shown in Fig 8 : namely, *PR*_*PCA*_ are reported as red stars in panels (c) and (g), while *D*_*PCA*_(*p*) as squares in panels (d) and (h) for *p* = 60% (black), 70% (blue) and 80% (green). A first extremely evident difference with SVC-based dimensionality is that the values of the PC-based dimensions are quite similar for the PR, while for the *D*_*PCA*_(*p*) the SY1 data are larger that the MA1 one, but at most by a factor of two. However, analogously to the SVCA we observe for both data sets a clear tendency of the PC-based dimensions to saturate to constant values for increasing number of neurons. This is consistent with what is reported in (Manley et al., 2024b) for *D*_PCA_(*p*). Analogous saturations have been shown for *PR*_PCA_ in (Dahmen et al., 2020) for neuropixels data obtained across different cortical areas in spontaneous behaving mice and in (Wang et al., 2025) for calcium imaging data from whole-brain of larval zebrafish hunting and in spontaneous conditions.

In order to characterize the evolution of the dimension estimates with the number of neurons we have performed a fitting to the data with the following simple expression

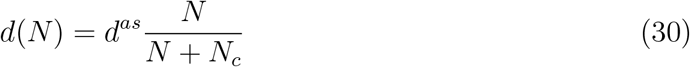

where *d*^*as*^ is the aymptotic value for the dimension and *N*_*c*_ sets the size above which the 50% of *d*^*as*^ is reached. The estimated asymptotic values for 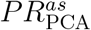 and 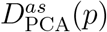 are reported in Table 3 and the fittings in Fig 8. In this case the values of 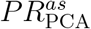 for SY1 and MA1 data are quite similar, only differing by a 13 %. For what concerns the 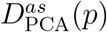, the asymptotic dimensions are larger for the SY1 data set with respect to the MA1 one, however the difference in this case are much smaller than in the SVC case, ranging from 40% to 55%.

Another interesting aspect is the fast convergence observable to the asymptotic values for the PRs with respect to the *D*_PCA_(*p*) estimates. Indeed, for the participation ratios 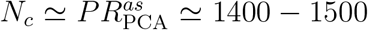, while for the *D*_PCA_(*p*) one has *N*_*c*_ ≃ 15000 − 20000 (*N*_*c*_ ≃ 8500) for SY1 (MA1) data.

For what concerns *PR*_PCA_ the scaling of this quantity with the number of neurons *N* have been derived theoretically in (Dahmen et al., 2020; Mazzucato et al., 2016). In particular, for *z*-scored data the participation ratio can be rewritten as follows

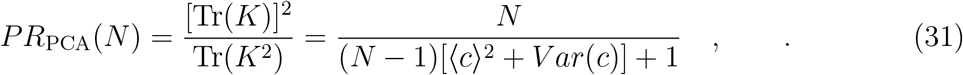

where *K* is the cross-correlation matrix as defined in (1), 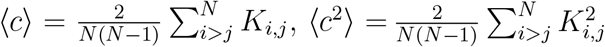 and *V ar*(*c*) = ⟨*c*^2^⟩ − ⟨*c*⟩^2^. Since we are considering *z*-scored data ⟨*c*⟩ coincides with the mean pair-wise correlation *ρ* and *V ar*(*c*) with the variance of the pair-wise correlations *V ar*(*ρ*) (Mazzucato et al., 2016).

The expression (31) coincides with (30) once set 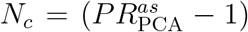 and 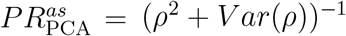. Indeed our data can be the fitted extremely well with expression (31), the results are essentially indistinguishable from the dashed red lines reported in Fig. 8 (c) and (g). The fitting procedure gives the following estimate for the only parameter entering in (31), namely *ρ*^2^ + *V ar*(*ρ*) = 0.000747 (0.000637) for SY1 (MA1) data. This estimate is in extremely good agreement with a direct evaluation of *ρ* and *V ar*(*ρ*) obtained from the data for different system sizes *N* ∈ [1024 : 32768]. In particular, we have observed a weak dependence on *N* of these two quantities and that the ratio (*ρ*^2^)*/*(*ρ*^2^ + *V ar*(*ρ*)) was of order 11% (1%) for SY1 (MA1) data, therefore *V ar*(*ρ*) *>> ρ*^2^ as already noticed in (Dahmen et al., 2020).

These indications suggest that in the cases that we have considered, 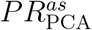 is essentially determined by the value of *V ar*(*ρ*) and not by the mean correlation *ρ*. Indeed, while 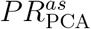 takes extremely similar values for SY1 and MA1 data sets (see Table 3), the corresponding correlations are quite different, namely *ρ* = 0.0090(1) (*ρ* = 0.0023(2)) for SY1 (MA1) data. The value found for the SY1 data set is consistent with those recently reported in (Pattadkal et al., 2026) for the primary visual cortex of marmorset and mice during spontaneous activity. The mean pair-wise correlation is definitely smaller for MA1 data, this can find its explanation in the fact that such data concerns neurons distributed across an entire hemisphere, that thus are expected to be less correlated.

## 4. Summary and Discussion

In this paper we have presented a detailed characterization of the Shared Variance Component Analysis (SVCA), a method specifically developed to perform dimensionality reduction of neural data originating from spontaneous activity (Stringer et al., 2019b). The aim of SVCA is to reveal the dimensions of the neural variance associated to reliable responses due to common hidden signals affecting the considered neural population.

As a first step, in order to fully understand potentialities and limitations of the SVCA methodology we compared the ability of SVCA and PCA in determining the dimensionality of a common underlying signal in a very simple example. In particular, we considered noisy synthetic data, where the reliable signal is a three dimensional chaotic drive (the Lorenz model) controlling linearly *N slaved variables* in presence of white or correlated noise. We have shown that SVCA is extremely effective in extracting the dimensionality of the reliable space from the data for any level of noise and for any correlation duration. On the contrary PCA is strongly affected by the noise and unable to separate the evolution of the common drive from the noisy data, whenever the noise amplitude is not negligible.

An important feature of the SVCA that we have identified by performing this analysis is that the reliable and total variances *S*_*k*_ and *S*_*k,tot*_ associated to the common underlying signal grow proportionally to the number *N* of slaved variables. This dependence allowed us to determine the *asymptotic* value of the fraction of reliable variance 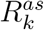 contained in the SVC dimensions and to determine that the linear dimensionality of the reliable space remains three for any system size. For the SVCA this result is confirmed by estimating also the linear *embedding* dimensionality (Jazayeri and Ostojic, 2021) by employing either the participation ratio *PR*_SVCA_ or *D*_SVCA_(*p*), which represents the minimal number of SVCs needed to recover the fraction *p* of the whole reliable variance.

This analysis has also revealed a drawback of the SVCA: the values of *S*_*k*_ for large *k* can become negative. This result is apparently contraddictory, since the quantity *S*_*k*_ represent variances, that in principle should be non-negative. This incongruence is due to the fact that the reliable variances (RVs) are obtained by a singular value decomposition of a cross-covariance matrix relative to the *test* period by employing a singular basis corresponding to a cross-covariance matrix estimated during a different period of time: the *training* one. However, we have shown that these spurious effects tend to disappear for sufficiently long test periods. Indeed, the sum of the negative RVs 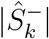 vanishes as 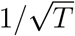, where is the duration of the test period.

The remaining of the paper has been devoted to the SVCA analysis of three large experimental neural recordings of the cortical dynamics of awake, spontaneously behaving mice. In particular, we have taken in consideration two-photon calcium-imaging data from the visual cortex V1 (Stringer et al., 2019b) (data set ST1, as defined in Table 1) and (Syeda et al., 2024) (data set SY1) and light bead microscopy data from the dorsal cortex (Manley et al., 2024b) (data set MA1).

Our first aim has been to reveal features of the SVC spectra that are common to all the considered experiments. A first important feature of these spectra, already pointed out in (Stringer et al., 2019a,b; Manley et al., 2024b), is that the RVs decay as a power-law *k*^−*α*^ with the SCV dimension *k*. This decay is observable in a range of SVC dimensions bounded by a maximal value *k*_*c*_(*N*) that grows with *N*. Moreover, the RVs in this range grow proportionally to *N*, thus strongly suggesting (in analogy with the synthetic Lorenz data) that this is the only part of the spectrum encoding for common reliable underlying signals.

Furthermore, by using a simple analytical argument we have demonstrated that the value of *k*_*c*_(*N*) grows as *N* ^1*/α*^ with the number of neurons *N*, with this theoretical scaling reproducing well an independent estimation reported by Manley et al. (2024b). This result suggests that the reliable spectra *S*_*k*_ will continue to extend the range where the powerlaw scaling is observable to smaller and smaller values of the variances by considering data sets with increasing number of neurons. This poses the question of whether these extremely small values contribute to the neural dimensionality of the reliable signal or not. Manley et al. (2024b) concluded that they are all relevant for the dimensionality estimation, down to values of order 10^−6^ (corresponding to the contribution of a single neuron over one million to the neural variance) and that therefore one has an unbounded growth of reliable dimensionality with the number of examined neurons.

We examined this growth of the reliable dimensionality from different point of views and with different methods, revealing that the dimension should first or late saturate to a maximal value. Let us first consider the definition of reliable dimensionality employed in (Stringer et al., 2019b; Manley et al., 2024b). This corresponds to the number of SVCs 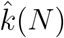 for which *R*_*k*_(*N*) ≥ Θ, where Θ is an arbitrary threshold. Since we have shown that an asymptotic profile for the fraction of reliable variance 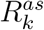 exists also for the experimental data, this implies that 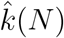 should saturate for sufficiently large *N* to a finite value 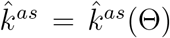 depending on the chosen threshold. Therefore the reliable dimensionality will not grow indefinitely with the number of neurons as we have shown by employing our analysis and some of the data reported in (Manley et al., 2024b). As an example, this procedure gives, by considering a threshold Θ = 90%, the following values for the dimensionality: 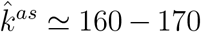, for ST1 data, 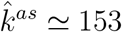 for SY1 data and 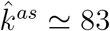 for MA1 data. On one side, these results are consistent with the ones reported in Stringer et al. (2019b), and also with the data in (Manley et al., 2024b), some of them being displayed in Fig. 7 (b). On another side, the results obtained for the ST1 and SY1 data sets corresponding to the same area of the cortex (visual cortex) are quite similar, while the MA1 data that encompasses an entire cortical hemispere has a definitely lower dimensionality. These differences are due to the smaller power-law decay exponents *α* measured for ST1 and SY1 data sets with respect to the MA1 one, as shown in Table 2. As previously pointed out in (Stringer et al., 2019a; Recanatesi et al., 2022) larger power-law decay exponents (with *α >* 1) are usually associated to lower dimensional signals.

As a second aspect, we have considered the embedding dimensionality of the reliable signal by estimating the corresponding linear dimensions *PR*_SVCA_ and *D*_SVCA_(*p*). These approaches, in order to obtain the linear dimensionality of the reliable space, give a different *weight* to each SVC dimension based on the reliable variance associated to such SVC. These are standard procedures to obtain neural dimensionality for PC-based analysis (Mazzucato et al., 2016; Engelken et al., 2023; Wang et al., 2025). For reliable spectra *S*_*k*_ decaying as a power-law we provide analytical expressions (27) and (28) for the participation ratio and for *D*_SVCA_(*p*), respectively. In particular, we have been able to provide analytical expressions for the dimensionalities for finite number *N* of neurons with power-law decaying spectra, while previously analogous analytical estimations have been reported only for infinite systems (Stringer et al., 2019a; Recanatesi et al., 2022).

The expressions in (27) and (28) depend only on the power-law decay exponent *α* of the spectra and they reproduce reasonably well the corresponding numerical estimations. The specific values for the considered data sets are reported in Figs 8 (a-b) and (e-f). In this case we limited the analysis to the SY1 and MA1 sets, neglecting ST1, since the data are quite similar to those reported in (Syeda et al., 2024) and limited to a smaller number of neurons. As a general feature we observe that *PR*_SVCA_ and *D*_SVCA_(*p*) are definitely larger for the SY1 data set than for the MA1 ones, almost one order of magnitude larger, and they display a clear tendency to saturate to some constant value for large *N*.

Another useful aspect of having an analytical formulation for the embedding dimensions is that we can easily derive their asymptotic values in the limit of an infinite number of neurons, with these asymptotic results reported in Eq. (29). As a general remark we observe that the asymptotic values are finite if *α >* 1. Therefore, for the considered data sets the values are always finite and the dimensionality will not grow indefinitely with *N*, but it will saturate. Moreover, the *PR*^*as*^ estimated with our approach reproduces quite well the exact expression (B.7) derived in (Recanatesi et al., 2022) as shown in Fig. B.3 in Appendix B. The specific values of the asymptotic 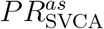 and 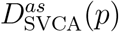 for the SY1 and MA1 data sets are reported in Table 3. The value found for SY1 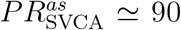 is of the order of the dimensionality reported in (Stringer et al., 2019b), we remember here that the SY1 data have been obtained in the same cortex area with an extremely similar methodology. In particular, Stringer et al. (2019b) affirms that SVCA has identified latent reliable signals encoded in at least 100 linear dimensions (possibly many more).

As a final aspect, to complete the analysis we have estimated PC-based embedding dimensions for the SY1 and MA1 data sets for growing number of neurons. The linear dimensions *PR*_PCA_ and *D*_PCA_(*p*) reveal a clear tendency to saturate, in agreement with the analysis reported in (Mazzucato et al., 2016; Dahmen et al., 2020; Manley et al., 2024b; Wang et al., 2025). However, the traditional PCA, without cross-validation to measure reliability as in SVCA, give a much larger estimates of the dimensionality of the neural activity, since now reliable and unreliable signals are not separated.

By limiting to the PR we have shown that the asymptotic values of PC-based dimensionality are controlled by the fluctuations of the pair-wise correlations, while are weakly influenced by the mean correlation values, in agreement with (Dahmen et al., 2020). In particular, SY1 and MA1 data show quite similar values for the variances of the correlations and therefore they also do for 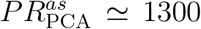. As we show in what follows, the asymptotic SVC-based estimations of the linear emedding dimensions are definitely smaller. In particular, we note that SY1 data have a dimensionality 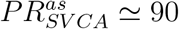, almost twenty times larger than that of MA1 data, namely 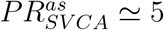. Therefore, on one side the PCA indicates that the overall neural activity for SY1 and MA1 data is described by almost the same number of PC dimensions, while on the other side the SVCA tells that the SVC dimensions encoding for the most part of the reliable variance are definitely fewer in the MA1 case with respect to the SY1 one. As already mentioned, this difference can be due to the origin of the data, while the SY1 data are taken by a restricted portion of the cortex (the visual cortex V1) the MA1 data refers to the entire left hemisphere. Indeed, this induces in the MA1 data a lower level of coherence (the mean pair-wise correllation is *ρ* ≃ 0.002) with respect to SY1 data (*ρ* ≃ 0.01) originating from the same cortical area. Therefore this leads to a reduced number of common reliable signals and at the same time to a larger amount of uncorrelated activity.

In summary, we have shown that the dimensionality of reliable neural signals associated to spontaneous activity is necessarily saturating with the number of considered neurons. This analysis has been performed by considering experimental data as well theoretical estimations for power-law decaying SVC spectra. As a general result the reliable dimensionality decreases for increasing values of the power-law exponent *α* and asymptotically (for *α >>* 1) only one reliable dimension will remain, as shown in Fig. B.3. Furthermore, we have presented a methodology to obtain asymptotic estimates of the reliable dimensionality based on limited samplings of the neuronal population, thus not requiring of a extremely large number of neurons to be achieved.

An interesting side result obtained with our analysis concerns the validity of the classical excitatory-inhibitory balance theory often employed to describe the asynchronous activity of the cortex (Van Vreeswijk and Sompolinsky, 1996; Renart et al., 2010). As already mentioned the mean *ρ* and the variance *V ar*(*ρ*) of the pair-wise correlations do not show any noticeable dependence on the number of neurons in our data sets, while accordingly to (Renart et al., 2010) they should both vanish as 1*/N*. This scaling would imply that the PC-based participation ratio (31) should grow proportionally to *N*, as previously reported for balanced neural networks in (Clark et al., 2023; Engelken et al., 2023). Instead, we observe a saturation of the participation ratio induced by the presence of small but not-vanishing pair-wise correlations (Mazzucato et al., 2016; Wang et al., 2025; Dahmen et al., 2020). Extremely weak correlations, of the same order of those here measured, have been recently shown to be also at the origin of the spiking variability observable in the cortex (Pattadkal et al., 2026). Therefore, our results and the ones reported in (Pattadkal et al., 2026) are challenging the classical balance theory.

A fundamental unsolved question is the origin of the power-law decay observable for the reliable variances as a function of the SVC dimension. This is a key aspect emerging from the cross-validation and that can be eventually related to the organization of neurons in spatial clusters as suggested in (Manley et al., 2024b), however devoted analysis are required in the future to fully capture the origin of this power-law decay that seems quite universal emerging both for spontaneous (Stringer et al., 2019b) and stimulus-evoked activity (Stringer et al., 2019a).

The SVCA is not limited to neuroscience data, it can find applications in other contexts. For example, SVCA can be useful to eliminate the irrelevant features present in huge data sets and to provide *reliable* data with optimal dimensionality, e.g. to avoid the well known overfitting problems in Machine Learning. Indeed, this could represent an interesting direction to investigate for future developments of the analysis here presented.

## CRediT authorship contribution statement

**Alejandro Carballosa**: conceptualization, data curation, formal analysis, investigation, methodology, software, validation, visualization, writing - original draft, review and editing. **Alessandro Torcini**: conceptualization, formal analysis, funding acquisition, investigation, methodology, project administration, resources, supervision, validation, visualization, writing - original draft, review and editing.

## Data availability

The synthetic data can be replicated following the models and parameters described in the methods. The experimental datasets considered in this work are openly available in their original repositories, see Stringer et al. (2018), Syeda et al. (2023), Manley et al. (2024a).

## Acknowledgments

The authors acknowledge useful discussions with Alberto Bacci, Rainer Engelken, Nina La Miciotta, Gianluigi Mongillo and are grateful to Carsen Stringer for providing some of the data. We are particularly in debt with Thomas Kreuz for a carefull reading of the manuscript before submission. A.C. and A.T. received financial support from the Labex MME-DII (Grant No. ANR-11-LABX-0023), from CY Generations with the project SLLOWBRAIN (Grant No. ANR-21-EXES-0008) and from FSM CY Initiative.

## Appendix A. Correlated Noise

One important assumption of SVCA is that the noise terms associated to single neurons are uncorrelated, this hypothesis allows for the cross-validation between train and test timepoints leading to the cancellation of the noise terms themselves. However, single neuron activity presents correlations of intrinsic fluctuations across cortical areas in behaving primates lasting hundreds of ms (Murray et al., 2014), moreover these timescales in presence of adaptive mechanisms can become of the order of tenths of seconds (La Camera et al., 2006).

Therefore, it is important to verify the validity of the above assumption by considering additive correlated noise (e.g. obtained from an Ornstein-Uhlenbeck (OU) process, see Methods) for increasing correlation times *τ >* 0. For this purpose we repeat here the same analyses performed for uncorrelated white noise in subsection (3.1). In particular, since we want to consider only the effect of the increased correlation time, we keep constant the variance of the OU process fixed at *V ar*(*ξ*) = 25 by adjusting the amplitude of the noise 𝒞 by following Eq. 19, see Methods for more details. In particular, the uncorrelated case (*τ* = 0) with the same variance of the noise would correspond to a noise-to-signal ratio value *ε* = 0.1.

As we can appreciate from Fig. A.1 (a) and (c) an increase on *τ* has no effect on the value of the first three variances *λ*_*k*_ and *S*_*k*_ obtained with PCA and SVCA, which remain equal to the value obtained for the white noise. The differences begin to emerge at *k >* 3, as shown in the insets of Fig. A.1 (a) and (c). In particular, for the PCA the spectrum at *k >* 3 does not satisfy anymore the expression (20): one has a strong increase of these variances with the correlation time *τ* with respect to the uncorrelated case. In terms of embedding dimensionality, we see that *τ* has no effect on the value of the participation ratio (*PR*_PCA_ = 3.38 − 3.36 in the studied range *τ* ∈ [0 : 10]) nor on *D*_PCA_(*p*) for *p* ≤ 80% (namely, *D*_PCA_(50%) = 2, *D*_PCA_(80%) = 3). A clear effect is evident only for *D*_PCA_(*p* = 95%), which decreases from a value of 214 at *τ* = 0 to 58 at *τ* = 10. This is due to the fact that at *τ* = 10 the variances *λ*_*k*_ are much larger between 3 *< k <* 100 than at *τ* = 0.

**Figure A.1:**
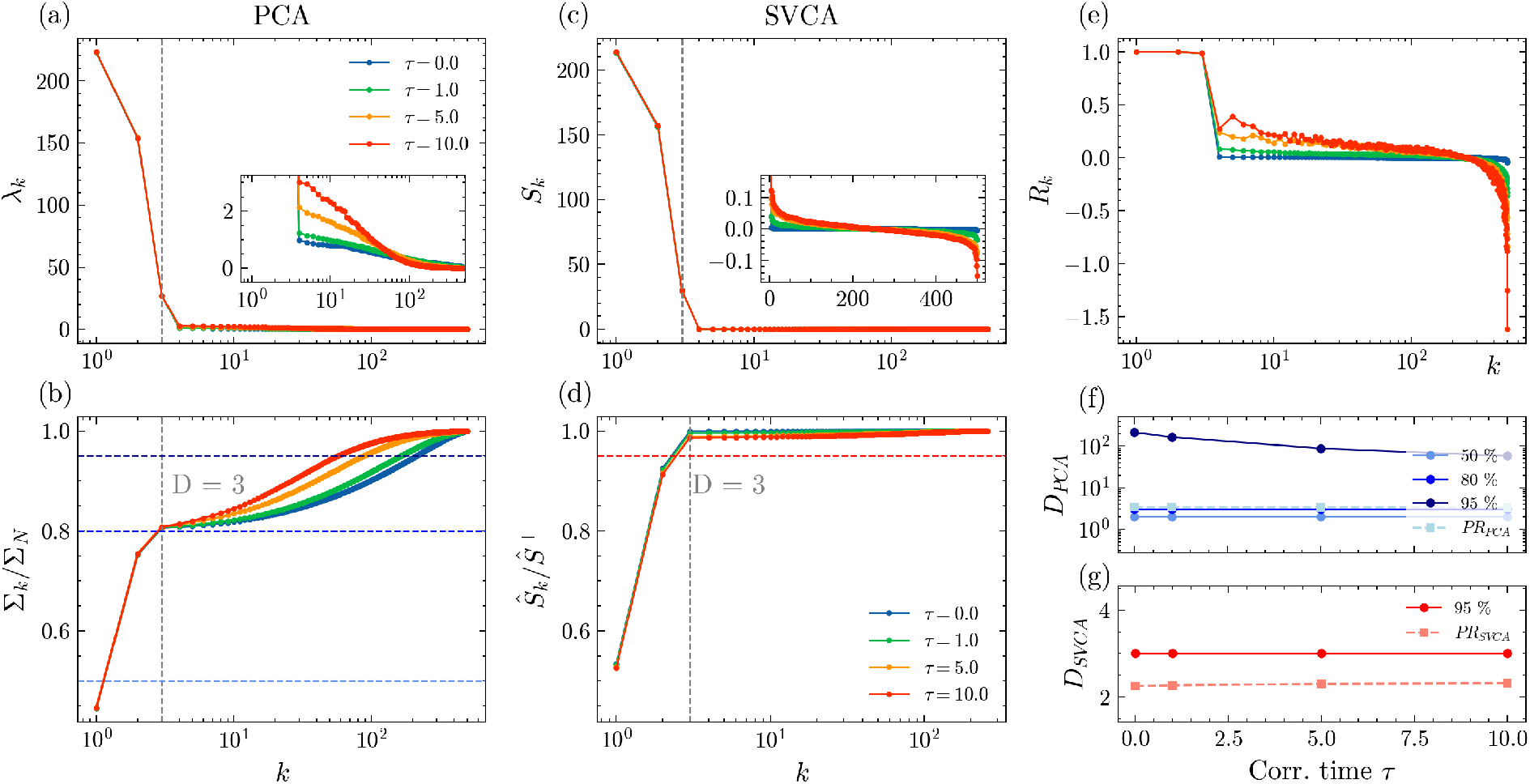
Lorenz system lifted onto a *N* = 500 dimensional space with correlated Ornstein-Uhlenbeck noise for different correlation times *τ* : namely, *τ* = 0 (i.e., white noise, blue symbols), *τ* = 1 (green symbols), *τ* = 5 (orange symbols) and *τ* = 10 (red symbols). (a) Spectrum of the PC eigenvalues *λ*_*k*_ versus their index *k*. In the inset an enlargement of the values for *k >* 3. (b) Normalized cumulative sum (variance) Σ_*k*_*/*Σ_*N*_ of the PC spectrum. (c) Spectrum of the RVs *S*_*k*_. In the inset an enlargement of the of the values for *k >* 3. (d) Normalized cumulative sum *Ŝ*_*k*_ */Ŝ*_*N*_ of the RVs. The vertical dashed line in (a-d) denotes the value of the dimension of the phase space of the Lorenz model *D* = 3 (e) Fraction of reliable variance contained in the *k*-th Shared Variance Component (SVC). (f) System dimensionality *D*_PCA_(*p*) estimated by employing PCA as in Eq. (13) for *p* = 50% (medium blue), 80% (blue) and 95% (dark blue) versus *τ*. The colors refer to the horizontal dashed lines in (b). In the same panel it is reported also the participatio ratio *PR*_SVCA_ (light blue) defined in Eq. (14) for the SVCA spectrum. (f) Dimensionality *D*_SVCA_(*p*) (red symbols) associated to the RVs for *p* = 0.95. In the panel it is also reported the *PR*_SVCA_ (orange symbols) estimated by employing Eq. (14) limited to the the positive part of the SVC spectrum *S*_*k*_. The number of sampling time points is *T* = 5 · 10^4^.

Furthermore, SVCA is still able to discern correctly the correlated noise as part of the unreliable signal for every value of *τ* considered. The spectrum associated to the noise (*k >* 3, inset of panel of Fig. A.1 (c)) is again symmetric around zero, obeys the extensive property with the system size *N* defined in Eq. (21). Moreover, the part of the RV spectrum *S*_*k*_ with *k >* 3 also decreases as 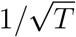 for increasing duration *T* of the test period analogously to the pure correlated noise signal reported in Fig. A.2. The main difference with respect to the uncorrelated case is that now the amplitude of the reliable variances *S*_*k*_ for *k >* 3 are definitely larger both for positive and negative values. In particular, the variances at *τ* = 10 are around one order of magnitude larger than those at *τ* = 0. This effect becomes dramatically evident when considering the fraction of reliable variances *R*_*k*_ reported in Fig. A.1. (e)).

In terms of embedding dimensionality, *τ* has also no effect due to the fact that the first three terms of the spectrum of the reliable variances take values definitely larger thna the rest of the spectrum. Indeed, we recover for any considered *τ* the same values obtained for the white noise, namely *D*_SVCA_(95%) = 3 and *PR*_SVCA_ ≃ 2.25 − 2.31.

**Figure A.2:**
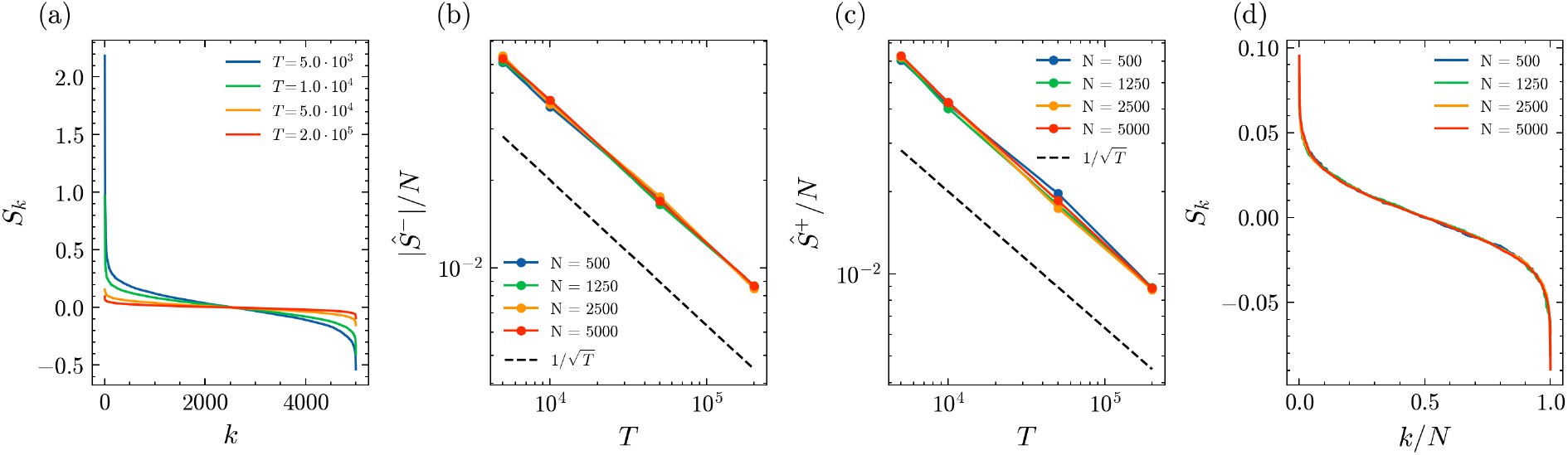
Correlated noise characterized in terms of the SVCA for correlation time *τ* = 1. (a) SVC spectra *S*_*k*_ measured for different number *T* of sampling data points for fixed system size *N* = 5000. (b) SVC spectra *S*_*k*_ versus *k/N* for different system sizes *N* ranging from 500 to 5000 for a number of timepoints *T* = 2 · 10^5^. The sum of the negative (positive) SVC eigenvalues |*Ŝ*^−^|*/N* (*Ŝ*^+^*/N*) is reported in panel (c) (panel (d)) rescaled with the system size *N* versus *T* for different system sizes. The dashed line denotes a decay 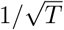. Here the variance of the OU process is fixed to *V ar*(*ξ*) = 1.

## Appendix B. Embedding dimensions for spectra decaying as a power-law

In this Appendix we will estimate analytically the expressions for the *D*(*p*) and the *PR* whenever a spectrum {*λ*_*k*_} (this can be either PCs, RVs or TVs) decays as a power law with the index *k* of the considered component. Therefore let us assume that

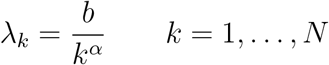

where *α >* 1 is the power law exponent and *b* a constant eventually dependent on the system size.

Let us first consider the estimation of the embedding dimension *D*_*p*_. Therefore we need to evaluate the sum Σ_*q*_, a reasonable approximation for finite *q* can be the following

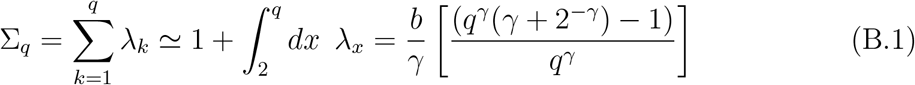

where *γ* = *α* − 1 and in order to estimate the sum we passed from discrete indeces for *k* ≥ 2 to continuous ones *x* ∈ [2, *q*].

From the definition of *D*(*p*) we have

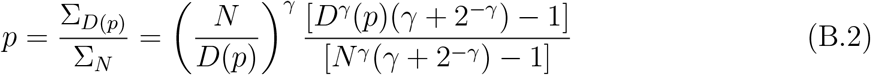

and finally

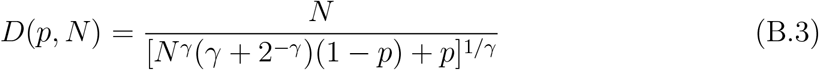

that is the expression Eq. (28) reported in the main tex. From this it is easy to obtain the expression reported in Eq. (29) for *D*^*as*^(*p*) in the limit *N* → ∞ and *α >* 1

In order to estimate the participation ratio we need to know also the sum of the squared eigenvalues, namely

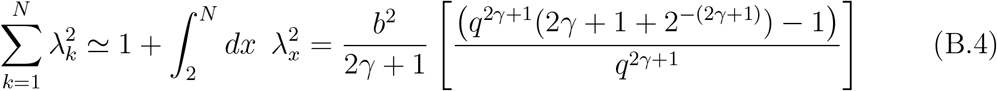

from the definition of the PR (14) we readily find the expression reported in the main text Eq. (27)

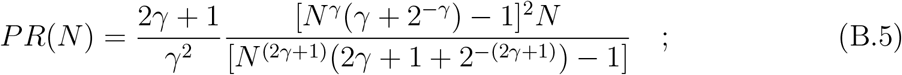

as well as *PR*^*as*^ (29) for *N* → ∞ and *α >* 1 (*γ >* 0).

As previously noticed in (Recanatesi et al., 2022) in the limit *N* → ∞ the sums Σ_*N*_ coincides with the Riemann zeta function *ζ*(*α*), apart a factor b, i.e.

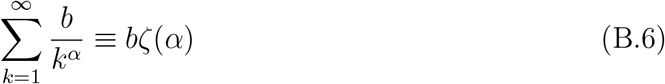

and *PR*^*as*^ has the following exact expression

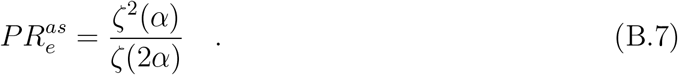

As shown in Fig. B.3 our estimation *PR*^*as*^ is in quite good agreement with the exact expression over the whole range of *α*. Furthermore, for very large *α* both *PR*^*as*^ and *D*^*as*^(*p* = 0.60) converge to one, consistently with what is observed for population responses in visual cortex for low dimensional stimuli (Stringer et al., 2019a). Furthermore, the asymptotic curves *PR*^*as*^ and *D*^*as*^(*p* = 0.60) respect the lower bound *d*_*m*_ given in (Stringer et al., 2019a) for the dimensionality of the linear space associated to a spectrum of the variances decaying as a power-law with an exponent *α*, namely *d*_*m*_ = 2*/*(*α* − 1). Apart for *D*^*as*^(*p* = 0.60) that for *α >* 1.5 is slightly lower than *d*_*m*_.

**Figure B.3:**
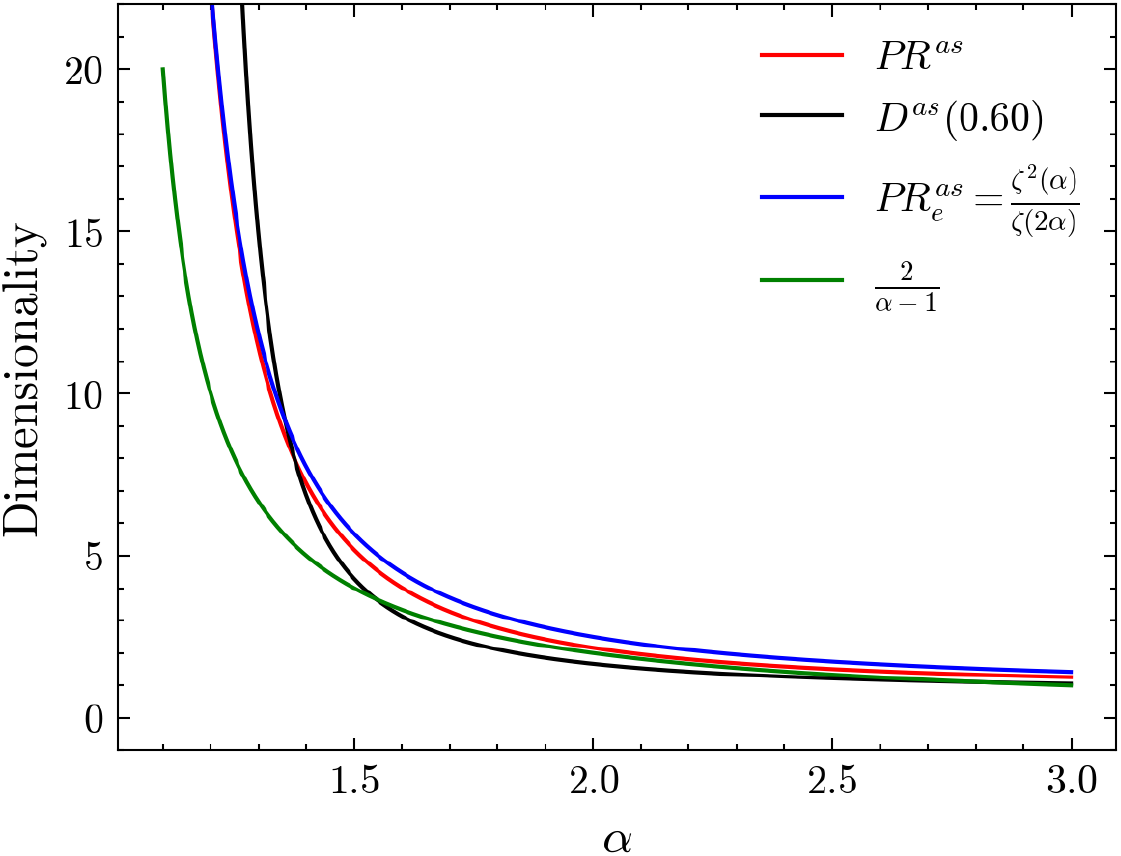
Dimensionality for a system with a spectrum of the variances decaying as a power-law with an exponent *α* versus *α*. The asymptotic values *PR*^*as*^ (red curve) and *D*^*as*^(*p* = 0.60) (black curve) in Eq. (29) are reported together with the exact value 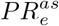 (B.7) (blue curve) and the lower bound *d*_*m*_ = 2*/*(*α* − 1) (green curve) derived in (Stringer et al., 2019a).

## Footnotes

1 Apart some small discrepancies observable at very large *N > T* : i.e. for *N >* 12596 (*N >* 4320) for the SY1 (MA1) data sets. This occurs whenever the number of recorded time points *T* becomes smaller than the system size *N* (thus resulting as the rank of the cross-covariance matrix since *r* = *T < N*). For the same reason in PCA whenever the PC-dimensions *k > T* the corresponding eigenvalues *λ*_*k*_ become exactly zero (Jolliffe and Cadima, 2016), in the SVCA this artifact introduces extra negative RVs, thus shifting the knee present in the data in the third column of Fig. 4 to the left and renders the scaling less evident.

